# Fine-scale population structure within and among Malagasy societies

**DOI:** 10.64898/2026.05.04.722645

**Authors:** Rindra Rakotoarivony, Evelyn J. Carter, Fernando Racimo, Denis Regnier, Jean Freddy Ranaivoarisoa, Mark Shriver, George Perry, Andrea Manica, Jason A. Hodgson

## Abstract

The population of Madagascar exhibits a globally unique combination of African and Asian genetic ancestries. Previous studies have described the admixture history of Madagascar at island-wide scales [1,2], but less focus has been paid to fine-scale population structure across the island. We present new genome-wide genetic data from 192 individuals sampled across five regions of Madagascar. We identify population structure at extremely fine spatial scales (∼10 km) among the Merina of the central highlands. By analysing subpopulations separately, we found one Merina group exhibited similarity to coastal populations in *f*_*4*_ ratios, estimated admixture dates, and pairwise *F*_*ST*_ distances, while another group was similar to other highland individuals in the same measures. This fine-scale substructure is likely associated with historical coastal-to-highland migration during the 18th and 19th centuries. In contrast, we also observe macro-scale structure in estimated timing of admixture across the island, with southeastern coastal groups exhibiting the earliest estimated admixture timings, and northern groups exhibiting the latest. This pattern corroborates previous results [1,2], and may suggest differing histories of admixture timing among Malagasy populations. Our results emphasise the importance of deep micro-geographic sampling to complement macro-scale analysis when characterising demographic history.

## Introduction

Situated at a juncture of historical trade routes along the Swahili coast and across the Indian Ocean, Madagascar is characterised by an unresolved history of migrations from Africa and Island Southeast Asia. Stone tools appear to indicate a human presence from 4,000 years BP [3], with possible evidence of human activity extending back to the early Holocene [4–6]. However, evidence of human activity is ephemeral until the first millennium CE when settlement sites become archaeologically visible [7–9]. Bantu-speaking east Africans from the Swahili coast [1,10] and Austronesian settlers from southeastern Borneo [11,12] arrived in Madagascar sometime between this first evidence of human activity and the first evidence of human occupation, beginning a cultural and ecological transformation of the island.

Gene flow between these disparate populations is reflected in the genomes of Malagasy individuals today, who inherit African and Asian genetic ancestry, with the proportion of each component varying between individuals [1,2,13,14]. Previous research has demonstrated that these genetic ancestry proportions are geographically heterogeneous, with coastal populations exhibiting higher proportions of African ancestry while highland populations exhibit greater proportions of Asian ancestry [2,10,15].

Bantu-speaking east-African individuals show the greatest similarity to the African component of Malagasy ancestry [1], and there has so far been no evidence of a more ancient African source contributing independently to this ancestry component [14], despite archaeological evidence indicating humans potentially reached Madagascar thousands of years earlier than previously assumed [3,4,6,16]. The publication of larger genotype datasets of Bantu African individuals [17], alongside new Malagasy data presented here, provide an opportunity to investigate the African component of Malagasy ancestry in greater detail.

Current models of the admixture process in Madagascar have lent support to the hypothesis that a single migration of very few Austronesian voyagers formed the basis of the Asian component of Malagasy ancestry [1,2,18]. Despite support for the hypothesis that the Malagasy population descends from a single wave of admixture [1,19], estimates of when this key admixture pulse began vary based on model assumptions, yielding dates between 22 and 30 generations ago [1,2,19]. Geographic variation in admixture timing across the island complicates this picture further [1,2,19]. The timing of genetic admixture in Madagascar is therefore still under scrutiny.

Systematic grid-based sampling combined with extensive genome-wide SNP genotyping provided a step change in understanding island-wide genetic diversity in Madagascar, by minimising geographic sampling bias and increasing the number of Malagasy samples available by ∼7 fold [2]. However, while extensive work has characterised the highland-to-coastal variation in genetic ancestry [2,10,15], less consideration has been paid to population structure at finer spatial scales. Dense sampling of individuals can build on the conclusions of macro-scale analyses by revealing important structure that may be missed when pooling samples for analysis at larger scales [20]. Given concentration on the question of when humans first reached Madagascar, fewer analyses have considered whether more localised post-admixture population structure can be observed across Madagascar.

Here, we analyse new genome-wide genotype data for 192 individuals sampled from five regions of Madagascar: Merina people from the Central Highlands (n = 36), Betsileo people from the Central Highlands (n = 38), Sakalava and Tsimihety people from the northwestern coast (n = 40), Betsimisaraka people from the eastern coast (n = 38), and people of varied ethnicity from the extreme north in the vicinity of Diego (n = 40). We ask: Which African populations are genetically most similar to the African component of Malagasy ancestry? Is there any observable population structure within Madagascar beyond the highland/coastal divide? When did the admixture event(s) that produced the modern Malagasy ancestry profile occur? And does the date of admixture vary between regions across Madagascar?

## Methods

Human genetic sampling was conducted under informed consent procedure approved by the Pennsylvania State University and the Madagascar Ministry of Health, and following community consultations with individual informed consent. Participants were recruited from five localities (Figure 1A), chosen to add to the geographic distribution of samples in Pierron et al. [14], and to include highland and coastal locations over a broad geographic area. Participants were asked their ethnicity and that of their grandparents. Saliva containing DNA was collected from each participant using Oragene collection tubes. DNA was extracted from saliva samples and purified using prepIT®•L2P, and DNA concentrations were standardised in preparation for genotyping using Nanodrop. 713,014 SNPs were typed using the Illumina HumanOmniExpress-24 v1.1 array.

**Figure 1:**
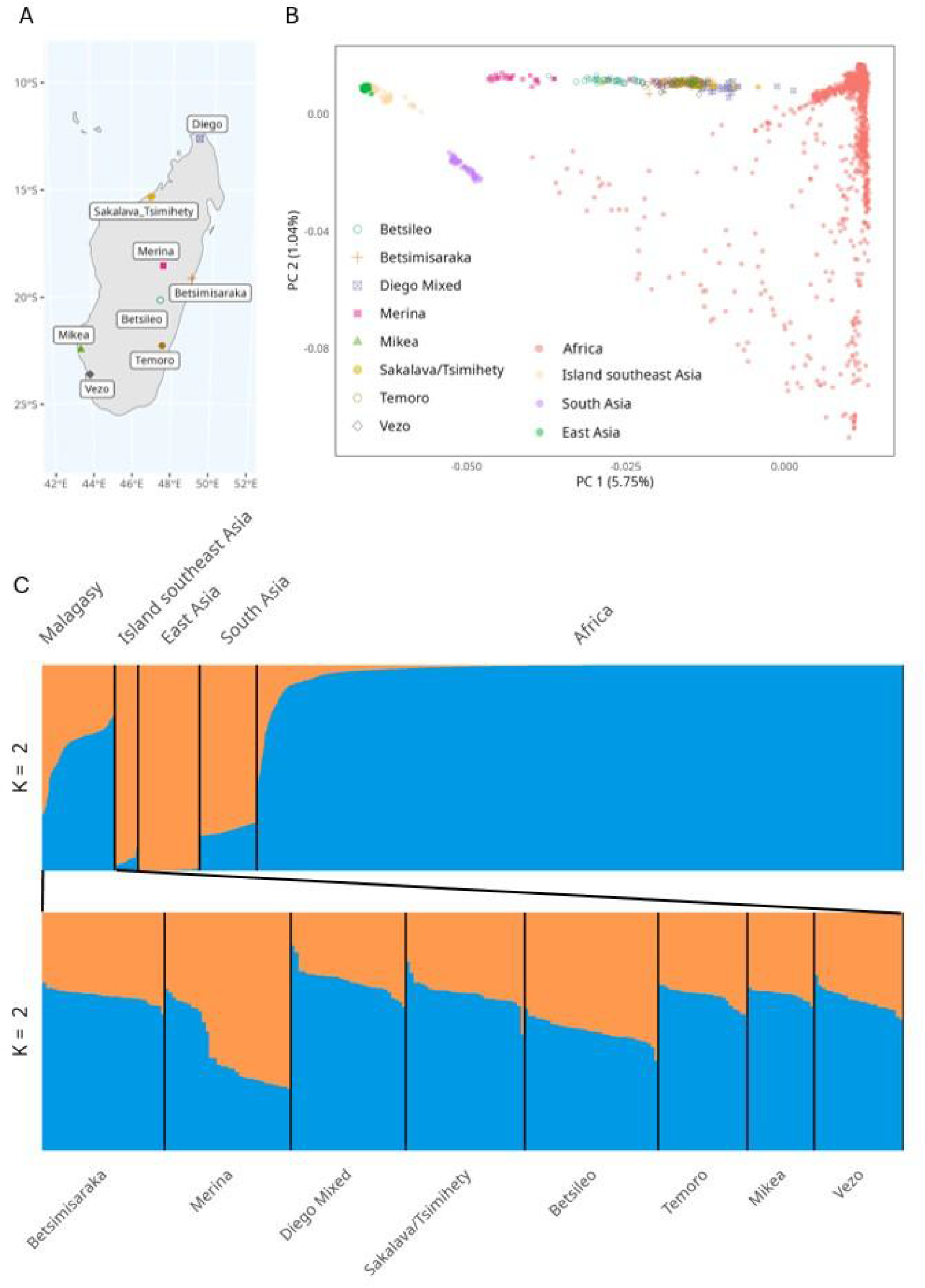
A) Map of Madagascar showing approximate sampling locations. Mikea, Temoro, and Vezo points are taken from [14]. B) Principal components analysis of Malagasy data (n = 232) combined with global datasets (n = 1754). C) ADMIXTURE results at K = 2 for Malagasy data compared to global populations. ‘Island Southeast Asia’ is composed of individuals from Borneo [11], and Sumba and Timor [22]. ‘East Asia’ is composed of Han Chinese South, China (CHS) and Kinh in Ho Chi Minh City, Vietnam (KHV) [21]. ‘South Asia’ is composed of Punjabi individuals sampled from Lahore, Pakistan (PJL) and Bengali individuals in Bangladesh (BEB) [21]. ‘Africa’ is composed of African individuals in the 1000 Genomes Project Consortium dataset [21], in addition to Bantu African individuals [17]. The 1000 Genomes Project populations include Esan in Nigeria (ESN), Gambian in Western Division Mandinka (GWD), Luhya in Webuye, Kenya (LWK), Mende in Sierra Leone (MSL), Yoruba in Ibadan, Nigeria (YRI).

Two main datasets were created, a Malagasy only dataset and a global comparative dataset. The Malagasy dataset contained our data merged with data collected by Pierron et al. [14] (Mikea; n = 21, Temoro; n = 24, Vezo; n = 24). The global dataset combined this set with southeast Asian and African populations in the 1000 Genomes Project Consortium dataset [21] (n = 882), additional African populations from Fortes-Lima et al. [17] (n = 1763), and populations from Borneo [11] (n = 39), Sumba (n = 30, subsampled randomly from the full dataset of 233 individuals) and Timor [22] (n = 7). Combining our data with Fortes-Lima et al.

[17] provided a larger comparative dataset of Bantu African populations than previous studies. Genome build of each dataset was verified using package `bigsnpr` [23] to ensure each dataset used build GRC37.

Data were merged in R using tidypopgen function `rbind` [24], which merges SNP datasets, resolves strand inconsistencies, and removes ambiguous SNPs. After merging, datasets were cleaned using tidypopgen [24] (Supplementary Information and Supplementary Table 2 contain quality control details). After filtering, the Malagasy dataset consisted of 233 individuals and 483,776 SNPs, and the global dataset consisted of 2,744 individuals and 215,593 SNPs. SNPs in linkage were removed by LD clumping [23], using tidypopgen. Those with an r^2^ value greater than 0.2 in a window of 100 kilobases were removed. After LD clumping, the Malagasy dataset consisted of 227,410 SNPs and the global dataset consisted of 198,196 SNPs.

Principal components analysis (PCA) was used to visualise Malagasy individuals compared to global comparative data. Non-Malagasy populations were randomly subsampled to a maximum of 30 individuals to avoid uneven sampling bias.

Unsupervised clustering using ADMIXTURE [25] was run on the cleaned global dataset using tidypopgen [24]. The algorithm was run for 9 values of K (2 to 10) with 5-fold cross validation (Supplementary Figure 3).

*f*_*2*_ statistics were calculated in tidypopgen [24], *f*_*3*_ and *f*_*4*_ ratios were then calculated in Admixtools2 from these *f*_*2*_ blocks [26]. Admixture *f*_*3*_ statistics were used in the configuration *f*_*3*_(Malagasy; Asian, African) to test whether the Malagasy could be modelled as an admixture between African and Asian references.

As the Merina exhibited high inter-individual variation in ADMIXTURE ancestry profiles, indicative of some form of population structure, we ran supervised ADMIXTURE [25] at K = 2 for all Malagasy populations [25] (Supplementary Figure 4). Borneo and Mozambique were used as references, as these populations gave the most negative *f*_*3*_ values when used as sources (i.e. they were the best putative sources in our dataset). Each population showed inter-individual variation in ancestry proportions (Supplementary Figure 5). However, the Merina exhibited a bimodal distribution (Supplementary Figure 6), as reported previously [13,15]. Merina with a high proportion of African ancestry and Merina with a high proportion of Asian ancestry were separated before calculating *f*_*4*_ ratios, running MALDER, and calculating pairwise *F*_*ST*_ (see details below for each analysis). Individuals were assigned to Merina A or Merina B based on their supervised ancestry profile at K = 2, with Merina A estimated to have >50% of ancestry derived from a population similar to Borneo individuals, and Merina B estimated to have >50% of ancestry derived from a population similar to Mozambique individuals. Splitting this sample reduced power, as only 11 individuals remained in Merina group B. However, this avoided violating model assumptions (e.g. the model of linkage employed by MALDER assumes no population substructure and panmixia) and enabled us to consider whether there may be differing demographic histories among the Merina.

*f*_*4*_ ratio statistics were used to examine the proportion of ancestry derived from Asian versus African sources for each population (Supplementary Table 6). To obtain these ratios, the global dataset was merged with a European reference (1000 Genomes GBR) and a chimpanzee reference (PanTro3.0, GenBank accession GCA_000001515.5) to act as outgroups. *f*_*4*_ ratios were calculated in the configuration *f*_*4*_(GBR,Chimpanzee;Malagasy,Borneo) / *f*_*4*_(GBR,Chimpanzee;Esan,Borneo) to estimate the proportion of African admixture and *f*_*4*_(GBR,Chimpanzee;Malagasy,Esan) / *f*_*4*_(GBR,Chimpanzee;Borneo,Esan) to estimate the proportion of Asian admixture. Standard errors were calculated using block-jackknife resampling (admixtools *f*_*4*_ ratio default).

MALDER [27,28] was used to estimate timing of admixture (Supplementary Table 7). Genetic distances were set using HapMap Phase II genetic map [29], mapped to build GRCH37. tidypopgen was used to create a subset of the full dataset containing only SNPs with known genetic map positions. This dataset was not pruned for linkage disequilibrium, and consisted of 425,904 SNPs. Standard errors were calculated through leave-one-chromosome-out jackknifing.

Pairwise *F*_*ST*_ between Malagasy populations was calculated in tidypopgen, using the method of Weir and Cockerham [30] (Supplementary Table 8).

## Results

We began by running a PCA using the new dataset combined with other Malagasy individuals and global comparative data. Malagasy individuals formed a cline between African populations and southeast Asian populations along the first principal component (Figure 1B, 5.75% of variance), representing the known geographic population structure between the highlands and the coast of Madagascar [2]. Individuals from Diego fell closer to African individuals, while some Merina individuals fell closer to southeast Asian individuals.

To examine variation among the Malagasy in greater detail, we ran unsupervised clustering algorithm ADMIXTURE [25]. Figure 1C shows ancestry profiles of Malagasy individuals compared to a global dataset at K = 2. East Asian, South Asian, and Island Southeast Asian individuals are primarily represented by the orange component, while African individuals are predominantly represented by the blue component. Malagasy individuals are a mixture of these, illustrating their admixed ancestry. There is considerable inter-individual variation within the Merina, where some individuals exhibit a much higher proportion of the Asian component than others (range 32% to 74%), consistent with the observation in Figure 1B that some Merina individuals fall closer to east Asian populations along PC1.

We then used *f*_*3*_ statistics to test whether Malagasy populations can be modelled as an admixture of African and Asian reference populations. Almost all combinations of references tested produced a negative *f*_*3*_ (6623 of 6776), indicating that the target could be modelled as an admixture between the references. The few cases that did not produce a negative *f*_*3*_ used either a mixed ancestry South African population as the African reference, or a Punjabi or Bengali population as the Asian reference, indicating these were not appropriate proxies for the sources of Malagasy admixture. Minimum *f*_*3*_ values were obtained from configurations using southern Borneo as the Asian source, and a Mozambique population as the African source (Supplementary Table 3). Figure 2A illustrates this, as the standard error of the *f*_*3*_ values using a Mozambique reference and the next best fitting African reference do not overlap. The same pattern is observed across each Malagasy population (Supplementary Figures 7-13). However, as z-scores are impacted by sample size, and Mozambique references had very low sample sizes (Mozambique_Ronga, n = 2; Mozambique_Chopi, n = 4; Mozambique_Tswa, n = 4), the Mozambique populations did not exhibit the lowest z-scores. Minimum z-scores were observed using DRC_Shi or DRC_Ngwi populations (n = 129 and n = 67 respectively) (Supplementary Table 4). Figure 2B shows the best fitting *f*_*3*_ values form a ‘corridor’, from central Africa across to the best fitting references on the east African coast, mirroring the trajectory of the Bantu expansion.

**Figure 2:**
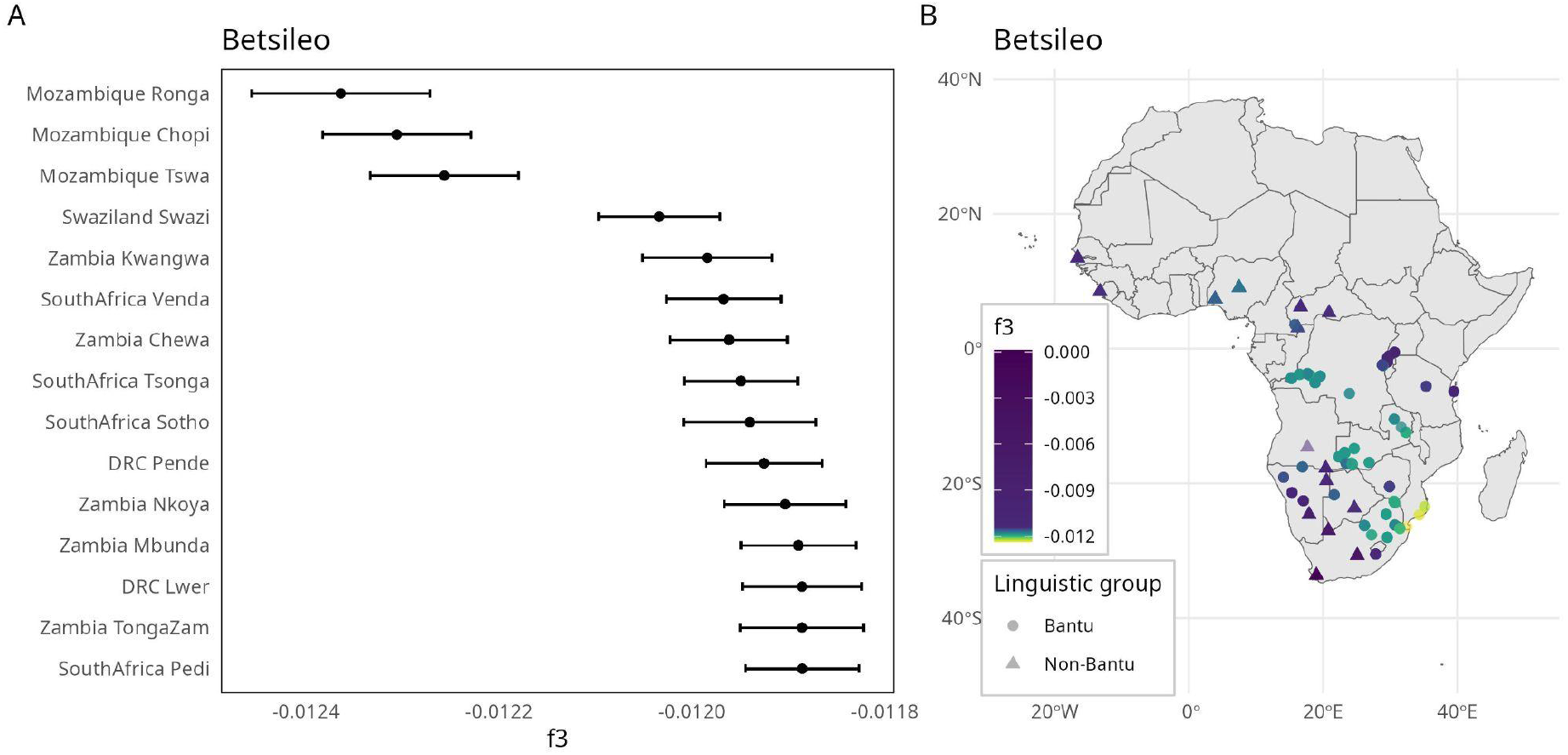
A) f_3_ statistic values across different African references for the Betsileo, showing the 15 lowest f_3_ estimates, with Borneo as the Asian reference. B) f_3_ statistic values across different African references, with Borneo as the Asian reference, plotted by location for populations where coordinates were available in metadata. Colour gradient was determined by taking the standard error of the minimum f3 value and multiplying this by 10. This scale was chosen to emphasise the very best fitting f3 values, and values outside this range are coloured dark blue.

The results of PCA and unsupervised clustering showed a high level of differentiation among the Merina. Individuals fell into two groups; those who clustered more closely with African individuals, and those who clustered more closely with Asian individuals. As this ancestry profile indicated population substructure, we used the supervised mode of ADMIXTURE [25], supplying individuals from Borneo and individuals from Mozambique as references, to infer the proportion of African versus Asian ancestry for all Malagasy individuals. Supervised ADMIXTURE illustrated a bimodal distribution of ancestry among the Merina, as previously identified by Hodgson [15]. We therefore estimated *f*_*4*_ ratios, pairwise *F*_*ST*_ distances, and admixture dates separately for Merina with a high proportion of Asian ancestry (>50%, labelled “A”), and Merina with a higher proportion of African ancestry (>50%, labelled “B”)(see Methods).

To explore the proportion of African to Asian ancestry across populations, *f*_*4*_ ratios were calculated. *f*_*4*_ ratios confirmed that Merina A had a lower proportion of African ancestry than Merina B (Merina A: alpha = 0.28, standard error = 0.01, z-score = 37.40, Merina B: alpha = 0.57, standard error = 0.01, z-score = 68.77) (Figure 3A). The lowest alpha value was observed among Merina A, while the Betsileo were intermediate between Merina A and the remaining populations (Betsileo: alpha = 0.46, standard error = 0.01, z-score = 76.94). Z-scores indicated these contributions were still significant for both populations. The highest alpha values were observed among Diego Mixed, and Betsimisaraka groups; these coastal populations exhibited the greatest proportion of African ancestry, and the corresponding z-scores indicated this contribution was significant (Diego Mixed: alpha = 0.62, standard error = 0.01, z-score = 110.60; Betsimisaraka: alpha = 0.61, standard error = 0.01, z-score = 107.40).

**Figure 3:**
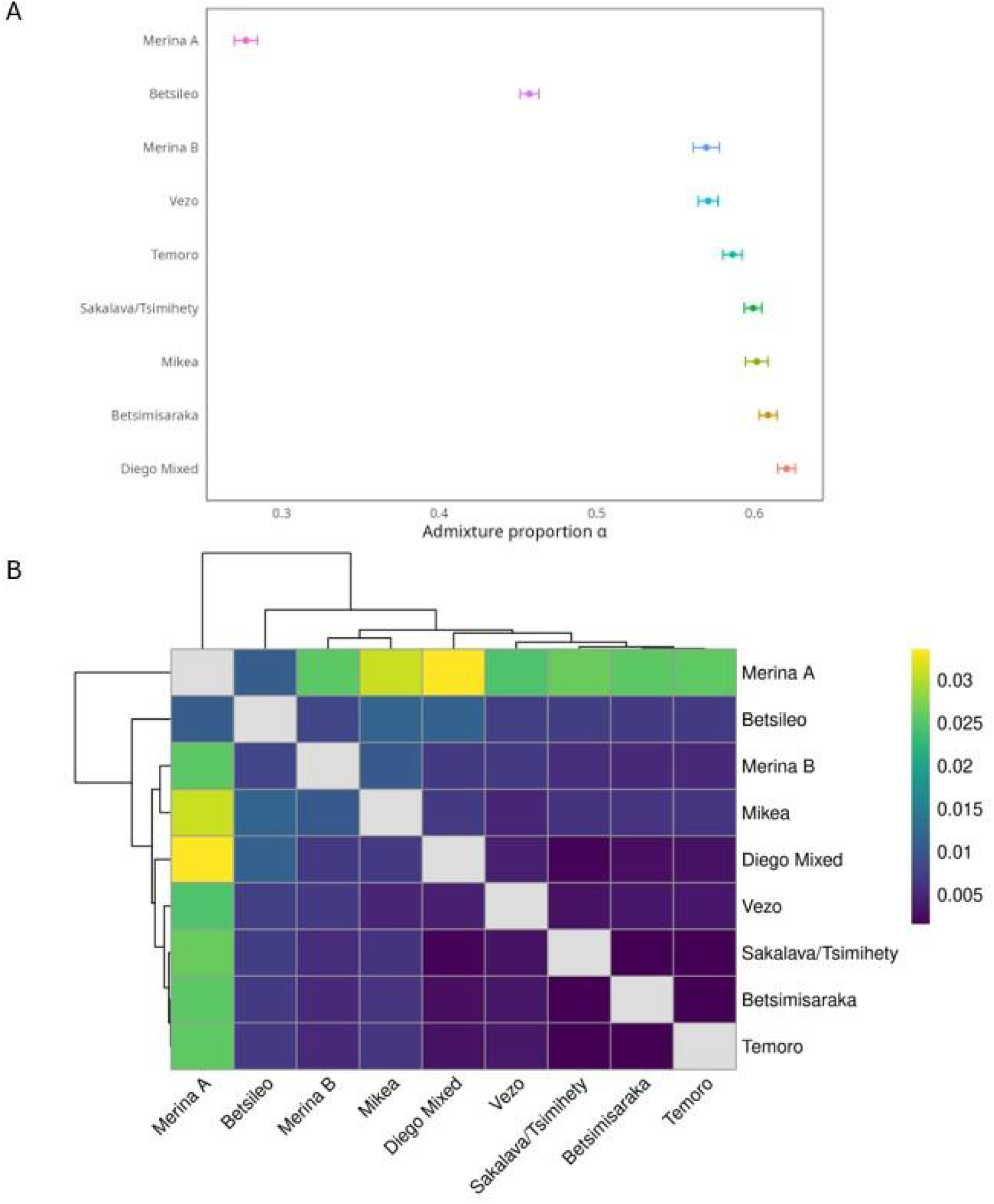
A) Proportion of admixture (α) from an African ancestry source into each Malagasy population. B) Pairwise F_ST_ estimates between Malagasy groups.

Pairwise *F*_*ST*_, was then used to examine differentiation between populations. The highest *F*_*ST*_ values were observed between Merina A individuals and all other Malagasy populations, with the highest values between Merina A and individuals from Diego (0.034) or Mikea individuals (0.031)(Figure 3B). The lowest *F*_*ST*_ values were observed between coastal group pairs (e.g. Diego Mixed and Sakalava/Tsimihety: 0.002, Supplementary Table 8). The highland Betsileo and Merina A outgroup all coastal populations, as expected given the known highland-to-coastal cline in genetic ancestry. However, Merina B fell within the branch containing all other coastal populations, and were clearly differentiated from Merina A (Figure 3B).

We used MALDER to explore the timing of admixture [28]. MALDER did not find support for multiple admixture events in any population. Assuming a generation time of 30 years [1,2], Figure 4 illustrates that the estimated timing of admixture fell between 600 and 900 years ago across Madagascar. Results for the Mikea, Vezo, and Temoro are consistent with previous estimates using ALDER [11,14] and MALDER [19]. Using reference populations with small sample sizes can generate noise in MALDER results [27]. However, our results show similar dates regardless of reference sample size. The estimated timing of admixture using a Mozambique reference with n = 4 did not differ significantly from the estimate using Yoruba (n = 104) or Esan (n = 98) (Supplementary Figure 14). The sources contributing to Malagasy admixture are deeply divergent, splitting from one another approximately 60,000 years ago [1], and admixture between them occurred recently (10’s of generations ago), meaning the references are not significantly diverged from the original admixing sources. Therefore, MALDER accurately infers admixture timing despite small sample sizes. We also observe a geographic pattern to admixture timing; the earliest dates are among the Temoro of the southeast and the Betsimisaraka in the east, while the latest are between Diego in the far north and the Vezo of the southwest (Figure 4). Interestingly, Merina A exhibited more recent dates of admixture than Merina B (Merina A = 21.86 +/- 0.88 generations; Merina B = 26.93 +/- 1.63 generations).

**Figure 4:**
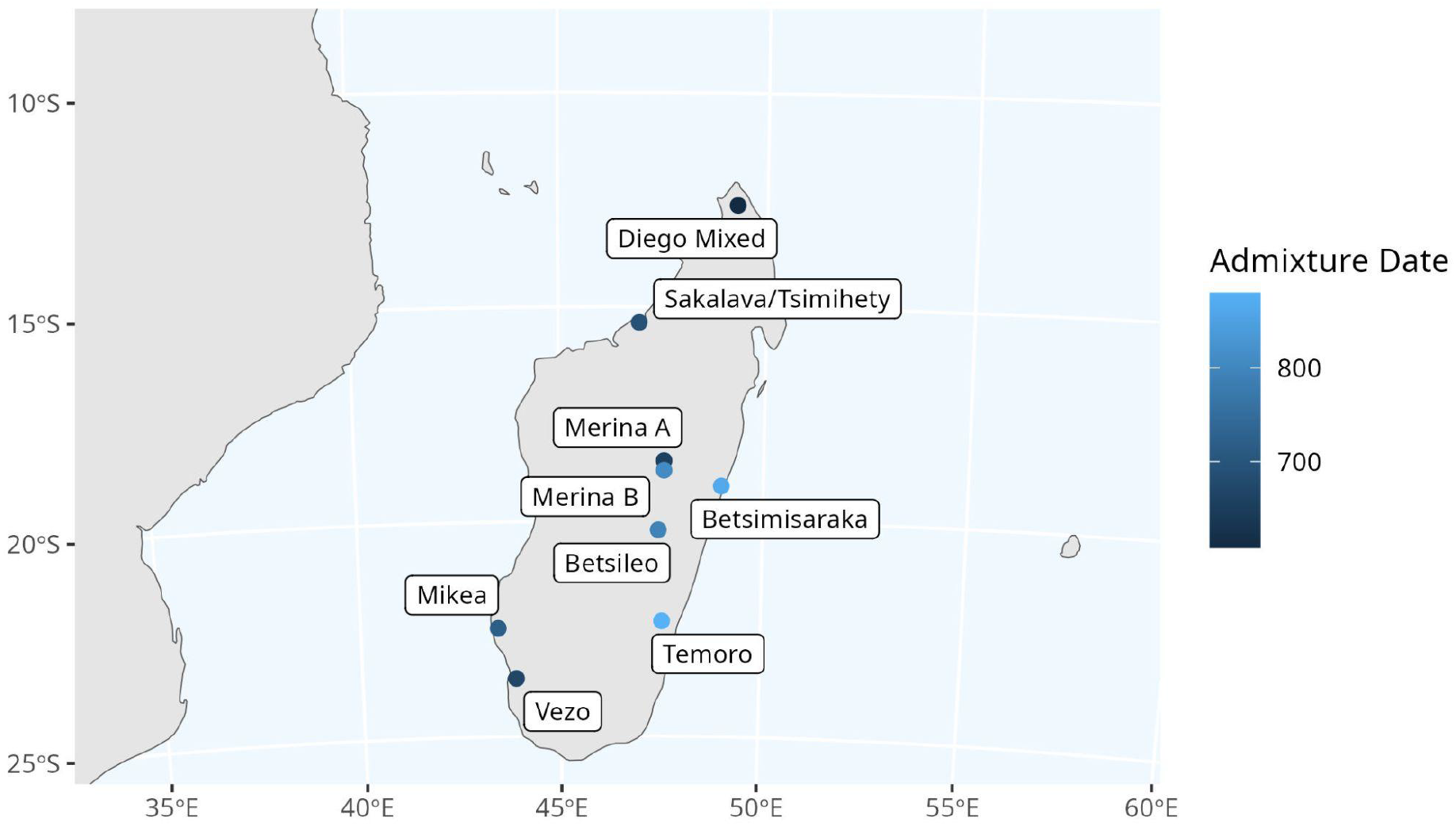
Map of Madagascar showing approximate sample locations and admixture dates as estimated by MALDER [28], with Mozambique as an African source population and Borneo as an Asian source population. Here, admixture date is calculated by multiplying the number of generations since admixture (the value given by MALDER) by an estimated generation time of 30 years.

## Discussion

We aimed to extend previous analysis by comparing Malagasy samples to a larger set of Bantu African reference populations [17] than previous studies. *f*_*3*_ statistics supported the conclusions of previous work, showing that Bantu-speaking African individuals from southern Mozambique (Ronga, Tswa, and Chopi) and individuals from southern Borneo (Banjar and Ngaju) best represent the ancestral sources that admixed to form Malagasy genetic ancestry [1,2,11]. Minimum *f*_*3*_ values were observed when using a Mozambique population as the African reference, however z-scores were lowest when using a Bantu-speaking African reference population with a larger sample size (DRC_Ngwi, n = 67; DRC_Shi, n = 129).

These results corroborate previous work showing Bantu-speaking eastern African populations to be the best fit for the African component of Malagasy ancestry using IBD [1].

Our results reject the hypothesis that multiple major waves of admixture produced the substrate of Malagasy genetic ancestry, corroborating previous findings [1,2,19]. However, admixture in Madagascar is relatively recent, meaning additional contributions from an African source some generations after admixture are unlikely to be reflected in the results of MALDER, especially if gene flow came from a population closely related to the original admixing source [27]. Two sources of periodic contact with Madagascar are the Comoros islands, and the east African coast. The Comoros islands have a similar demographic history to Madagascar, with present-day populations also being an admixture of African and Island Southeast Asian populations [19,31]. Archaeological [32] and anthropological [33–35] evidence attests to post-admixture migration between the islands, yet previous MALDER analysis obtained inconclusive results respecting Comorian admixture into the Malagasy [19]. Post-admixture contact with the east African coast is also evidenced, first by archaeology [9,32], and later by historical records of forced transport of enslaved people to Madagascar [32,34–38]. Therefore, while MALDER rejects a model of multiple waves of admixture, post-admixture gene flow may have impacted Malagasy genetic ancestry.

Continued contact with Africa also obfuscates the interpretation of ‘earlier’ and ‘later’ admixture events across the island. MALDER indicated that southeastern, eastern, and southern central highland populations have the earliest inferred dates of admixture (Temoro = 29.35 +/- 0.87 generations; Betsimisaraka = 28.84 +/- 0.92 generations; Betsileo = 26.44 +/- 0.82 generations), while the latest dates of admixture were observed in northern (Diego Mixed = 20.26 +/- 0.64 generations) and southwestern (Vezo = 22.26 +/- 1.24 generations and Mikea = 23.99 +/- 1.55 generations) populations. Pierron et al. [1] and Alva et al. [2] found the same geographic pattern, with the earliest dates occurring in eastern Malagasy genetic clusters (800 +/- 25 years BP [2]; 29.00 +/- 1.11 generations [1]) and the latest among northern clusters (665 +/- 19 years BP [2]; 19.99 +/- 0.95 generations [1]). The ‘later’ dates observed in populations with a greater proportion of African ancestry could reflect additional gene flow after initial admixture, for example, in the ‘Diego Mixed’ group who clustered close to modern Bantu African populations in PCA space (Figure 1B), exhibited the greatest proportion of African ancestry (Figure 3A), and showed the latest date of admixture (Diego Mixed = 20.26 +/- 0.64). While a single-wave model of admixture is well supported, the reality of continued gene flow between Madagascar and Africa is more complex.

Previous work based on historical and comparative linguistics has advanced the hypothesis that the Asian ancestors of the Malagasy first landed in the southeast of the island [39,40]. Although the earliest admixture dates are observed among individuals currently residing in the east and southeast of Madagascar, these cannot be used to support the hypothesis of an eastern ‘landing’. It should not be assumed that the location of present day populations reflects the geographic distribution of their ancestors, especially as historical records and oral histories attest to substantial internal migration in the period between admixture and the present day [41]. For example, Temoro (Antemoro) individuals from the southeast exhibit the earliest admixture date (Temoro = 29.35 +/- 0.87 generations), yet oral histories describe that ancestors of the Temoro migrated from the region of Vohémar on the northeast coast to establish the Temoro kingdom in the southeast at the end of the 15th century [42]. This introduces the possibility that an already admixed Temoro population arrived in the southeast and subsequently mixed with local people, exemplifying that conclusions about where admixture began or where the first Asian ancestors of the Malagasy landed cannot be drawn based on admixture dates alone.

Previous work has shown that coastal populations exhibit higher proportions of African ancestry while highland populations exhibit higher proportions of Asian ancestry [2,10]. Our analysis extended this work by considering whether population structure can be observed within or between Malagasy societies. ADMIXTURE profiles showed that coastal populations appeared more similar in their genetic profile to African populations than the highland Merina and Betsileo, and *f*_*4*_ ratio results confirmed that African ancestry proportion for each coastal population was greater than 50%. The central highland Betsileo exhibited less African ancestry than coastal groups (Figure 3A), as expected. However, the Merina of the central highlands had high inter-individual variation in African ancestry proportion (Figure 1C), corroborating previous findings [13,15]. Given this population structure, we split the Merina sample into two groups (see Methods), and *f*_*4*_ ratio results showed Merina A had a much lower proportion of African ancestry than Merina B (Merina A: alpha = 0.28, standard error = 0.01, z-score = 37.40, Merina B: 0.57, standard error = 0.01, z-score = 68.77), whose *f*_*4*_ ratio results appeared more similar to coastal populations (Figure 3A). Merina A individuals exhibited later dates of admixture (Figure 4), and higher values of pairwise *F*_*ST*_ distance to other populations (Figure 3B). In contrast, Merina B displayed admixture dates more similar to the Betsileo than to Merina A, and exhibited low pairwise *F*_*ST*_ distances to coastal populations (Figure 3B).

There are multiple possible causes for the observed population structure among Merina individuals. As Merina B individuals cluster closely with coastal populations in PC space, have ancestry proportions more similar to coastal groups, and exhibit low pairwise *F*_*ST*_ distances to coastal Malagasy groups, it is possible that recent migration generated this population structure. However, ethnographic accounts and historical records offer an alternative hypothesis. The Merina of Madagascar today inhabit a region, Imerina, where an initially small polity expanded to a Kingdom ruling over much of Madagascar from the 18th to 19th centuries [43]. During this expansion, the Merina Kingdom enslaved non-Merina Malagasy peoples and peoples from the east African coast [35,38,44], and transported many of them to the capital, Antananarivo, and Imerina. This transformed the region’s population, with some estimating that, at the time of abolition, enslaved people comprised 25% [44] to 50% [45] of the population in and around the capital. As a significant proportion of these enslaved people came from coastal Madagascar, from the northern and southern interiors, or from the east African coast itself, this movement broadened the spectrum of African genetic ancestry in Imerina. Before and after abolition, Merina social norms prohibited free people and their descendants from marrying enslaved people and their descendants [35]. The bimodal population structure observed among the Merina may, therefore, result from forced displacement followed by social norms precluding marriages between those with a greater proportion of Asian ancestry and those with a greater proportion of African ancestry. This may also impact the interpretation of estimated admixture timing. Merina A individuals exhibited a ‘later’ date of admixture, however, non-random mating according to genetic ancestry reduces the decay in linkage disequilibrium expected by MALDER, making admixture appear more recent [46]. Estimated admixture dates may therefore reflect the differing demographic histories of the Merina groups, and the impact of non-random mating.

Between 600 and 900 years ago, admixture took place between a population related to individuals from modern-day Mozambique, and a population related to individuals from modern-day Borneo, to establish the spectrum of genetic ancestry observed among the Malagasy today. Subsequent gene flow between continental Africa and Madagascar may have augmented the proportion of African ancestry across Madagascar since this initial admixture. However, we find no genetic signal that secondary ‘waves’ of migration contributed to the Malagasy genetic profile. Our analysis also revealed significant population structure at micro-geographic scale among the Merina of the central highlands. Previous grid-based sampling of individuals across Madagascar has transformed our understanding of Malagasy genetic ancestry [1,2], yet sparse sampling can miss fine-scale genetic structure [47], and aggregating data for analysis can obfuscate local-scale patterns of variation [20]. Fine-scale population structure can therefore be easily missed, but may reflect important aspects of population history. The population substructure we observe among the Merina, which has not been observed in previous studies, may be partly explained by the demographic impact of enslavement and slave trading within Madagascar. Future work should examine population substructure in greater detail to test this hypothesis, and to consider how more recent historical migrations have impacted Malagasy genetic ancestry.

## Acknowledgements

We would like to thank all the Malagasy people who participated in this research.

E.J.C. is supported by a Cambridge Philosophical Society Sedgwick studentship.

## Author contributions

R.R., J.F.R., J.A.H., M.S., and G.P. designed the study and performed data collection. R.R. and G.P. performed sample preparation for genotyping. F.R. and R.R. conducted preliminary data analysis. E.J.C., F.R., R.R., A.M., and J.A.H. contributed to the conceptualization of the data analysis. E.J.C. developed final analysis scripts. E.J.C. wrote the manuscript, with input from D.R., R.R., J.A.H., and A.M.. All coauthors reviewed the manuscript and provided critical feedback and suggestions. R.R and E.J.C contributed equally to this work.

## Ethical Approval

Ethical approval for collection of this dataset was obtained from the Ministère de la Santé Publique, Comité D’Éthique (Madagascar Ministry of Health, Ethics Committee) and from The Pennsylvania State University Institutional Review Board prior to data collection in 2015. Ethical approval for analysis of this dataset, in full accordance with the consent provided by participants when they entered the study, was granted by Anglia Ruskin University Faculty of Science and Engineering Biomedical Science School Research Ethics Panel and the University of Cambridge Council of the School of the Biological Sciences Human Biology Research Ethics Committee.

## Competing Interests

The authors declare no competing interests.

## Data accessibility

Malagasy genotype data generated by the current study will be deposited to the European Genome-Phenome Archive upon publication.

Genotype data for previously published Malagasy individuals can be accessed through the NCBI Gene Expression Omnibus (GEO) repository (accession number: GSE53445). [14], or are available in plink binary format on Zenodo at https://zenodo.org/records/17131856.

Genotype data from Bantu African individuals used in this study can be accessed via The European Genome-Phenome Archive at the European Bioinformatics Institute (accession number: EGAS50000000006). [17] Access to this dataset was granted by AfricanNeo Data Access Committee EGAC00001003398 on 18/08/2025.

Genotype data for individuals from Borneo can be accessed via The European Genome-Phenome Archive at the European Bioinformatics Institute (accession number: EGAD00010000944). [11]

Genotype data for individuals from the islands of Sumba and Timor are publicly available through the NCBI Gene Expression Omnibus GEO repository (accession number: GSE76645). [22]

## Notes

### Competing Interest Statement

The authors have declared no competing interest.

