## Supplementary Materials for "Fine-scale population structure within and among Malagasy societies"

### Cleaning and Merging Datasets

#### Supplementary

##### Data cleaning

Only autosomal SNPs were analysed. SNPs with a missingness rate above 0.05 were removed. Individuals with a genotype call rate less than 0.9 (missingness exceeding 0.1) were removed. SNPs with minor allele frequency below 0.025 were removed. Hardy-Weinberg equilibrium p-values were calculated for each SNP within each population using the tidypopgen function qc_report_loci, which provides a Bonferroni correction given the number of populations analysed. SNPs with a p-value below 0.01 in at least one population after correction were removed. Close relatives were removed using the KING robust kinship coefficient Manichaikul et al. (2010), implemented in tidypopgen. One individual was removed from each pair with a kinship coefficient greater than 0.0884, equivalent to removing relatives up to the second degree.

##### $f_4$ ratio

To obtain a chimpanzee reference, SNPs in our dataset were lifted to PanTro3.0 (GenBank accession GCA_000001515.5) from hg19 using liftOver overchain file https://hgdownload.soe.ucsc.edu/goldenPath/hg19/liftOver/hg19ToPanTro5.over.chain.gz from UCSC Genome Browser Kent et al. (2002), using bigsnpr function snp_modifyBuild() Privé et al. (2018). The chimpanzee reference for each SNP was obtained using a custom R function to query the Ensembl API (GenBank accession GCA_000001515.5, Ensembl release 115, accessed via Ensembl API https://rest.ensembl.org/documentation/info/sequence_region on 17-12-2025). This procedure retained 77993 SNPs for $f_4$ ratio calculation. Standard errors were calculated using block-jackknife resampling (admixtools $f_4$ ratio default). Supplementary Table 5 contains full $f_4$ ratio output.

##### MALDER

MALDER was run using the default -binsize of 5 * 10-4 Morgans (describing the size of the bins that SNPs are split up into. Instead of directly calculating distance between each SNP pair, MALDER uses the distance between the bins of each SNP pair.), and the –mindis parameter (minimum genetic distance to begin LD decay curve fitting) set to 0.005 cM.

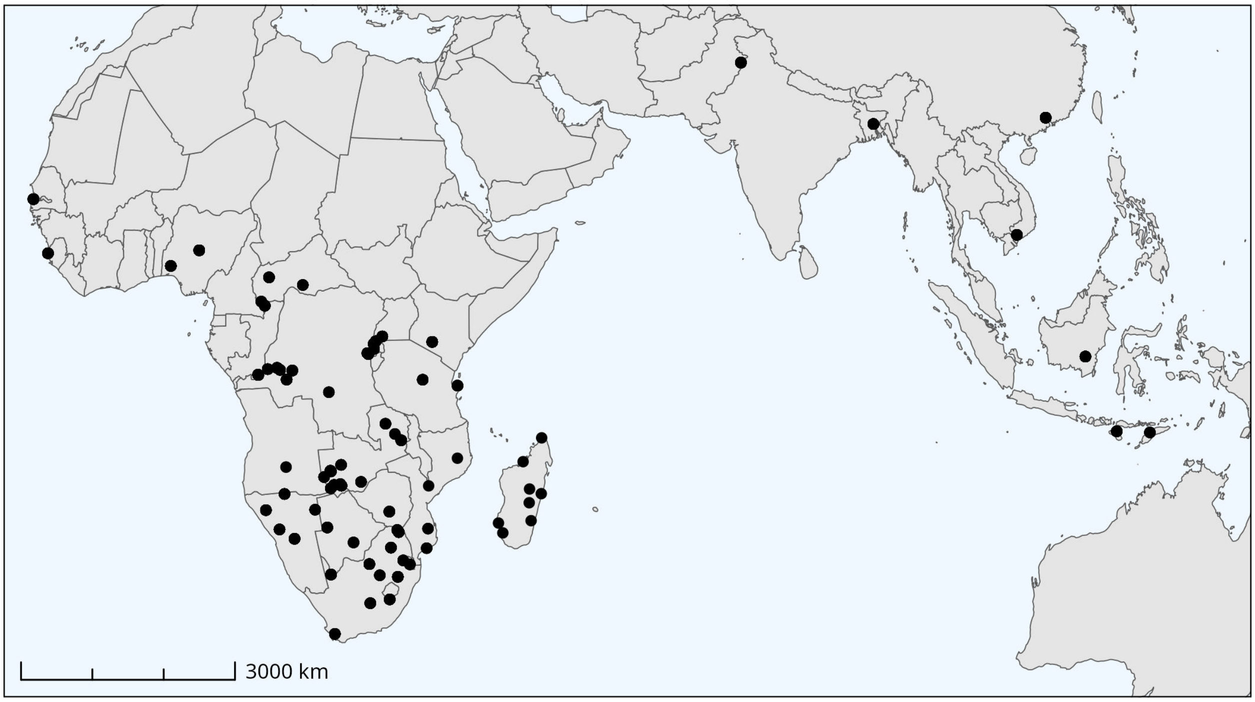


Figure 1: Map of reference population locations for all populations where coordinates were available in metadata.

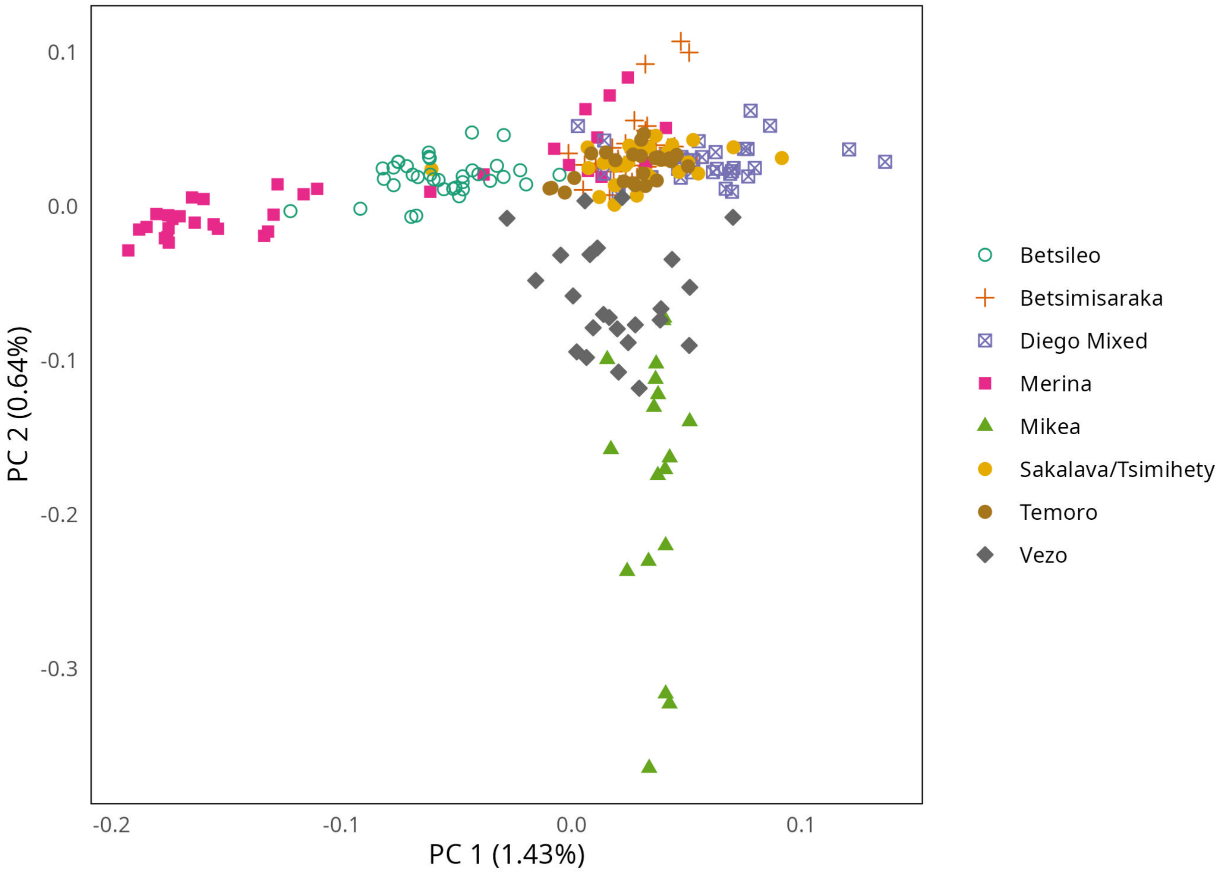


Figure 2: Principal components analysis of Malagasy data (n = 167), combined with data from Pierron et al. (2014) (Mikea, Vezo, and Temoro data points, n = 69). Colour and shape indicates self-identified ethnicity. 'Diego Mixed' individuals identified as Anjoaty, Antakarana, Sakalava Tavaratra, and Sakalava ethnicities.

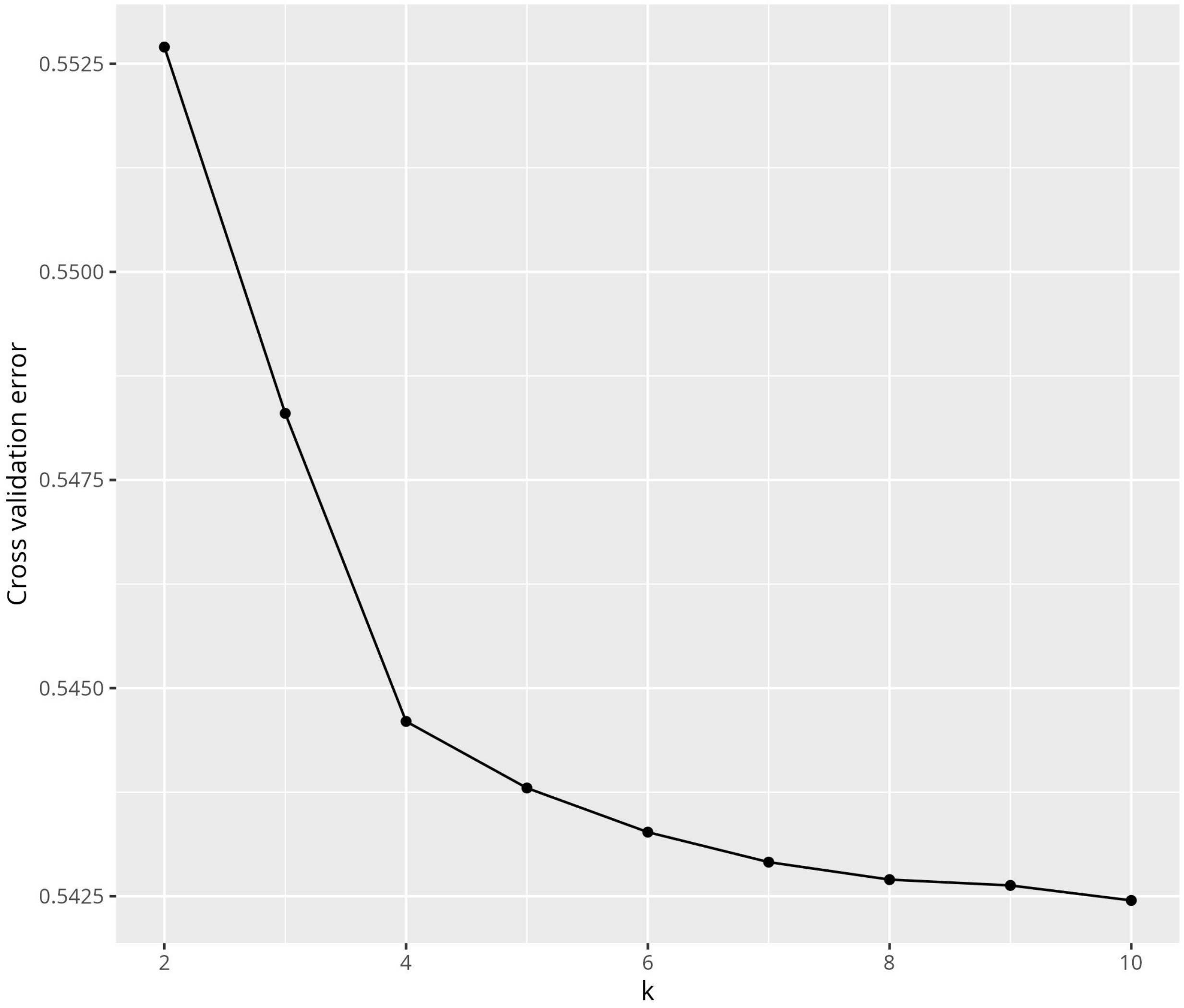


Figure 3: Cross-validation error for admixture.

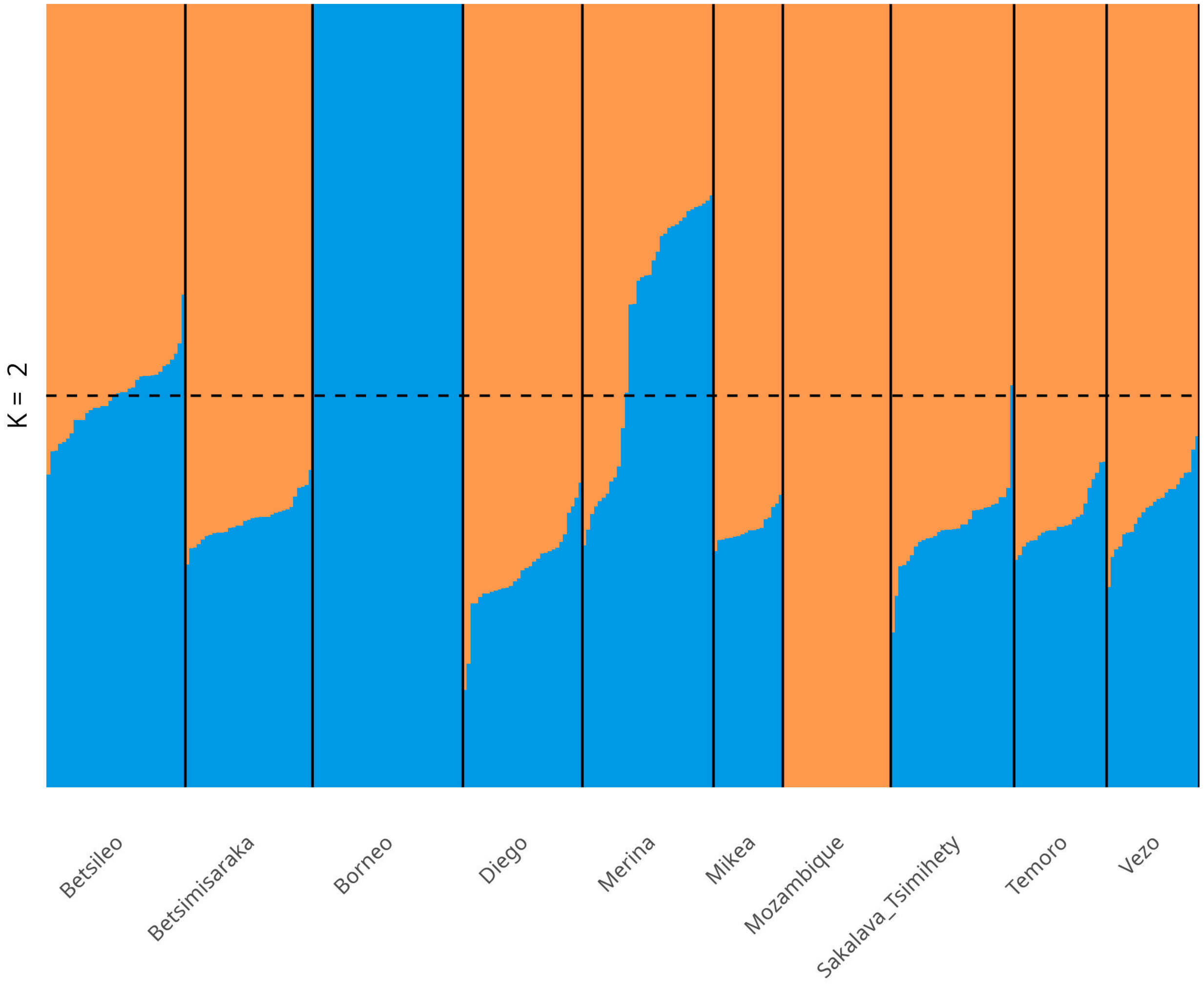


Figure 4: Supervised admixture at K = 2 for Malagasy compared to reference populations. Individuals from Borneo from Brucato et al. (2016), and individuals from Mozambique populations from Fortes-Lima et al., (2024). Mozambique individuals are labelled together, but this group contains individuals given the following population labels: "Mozambique_Nhungwe", "Mozambique_Tsonga", "Mozambique_Ronga", "Mozambique_TongaMoz", "Mozambique_Chuwabu", "Mozambique_Tswa", "Mozambique_Chopi", "Mozambique_Sena", "Mozambique_Makhuwa".

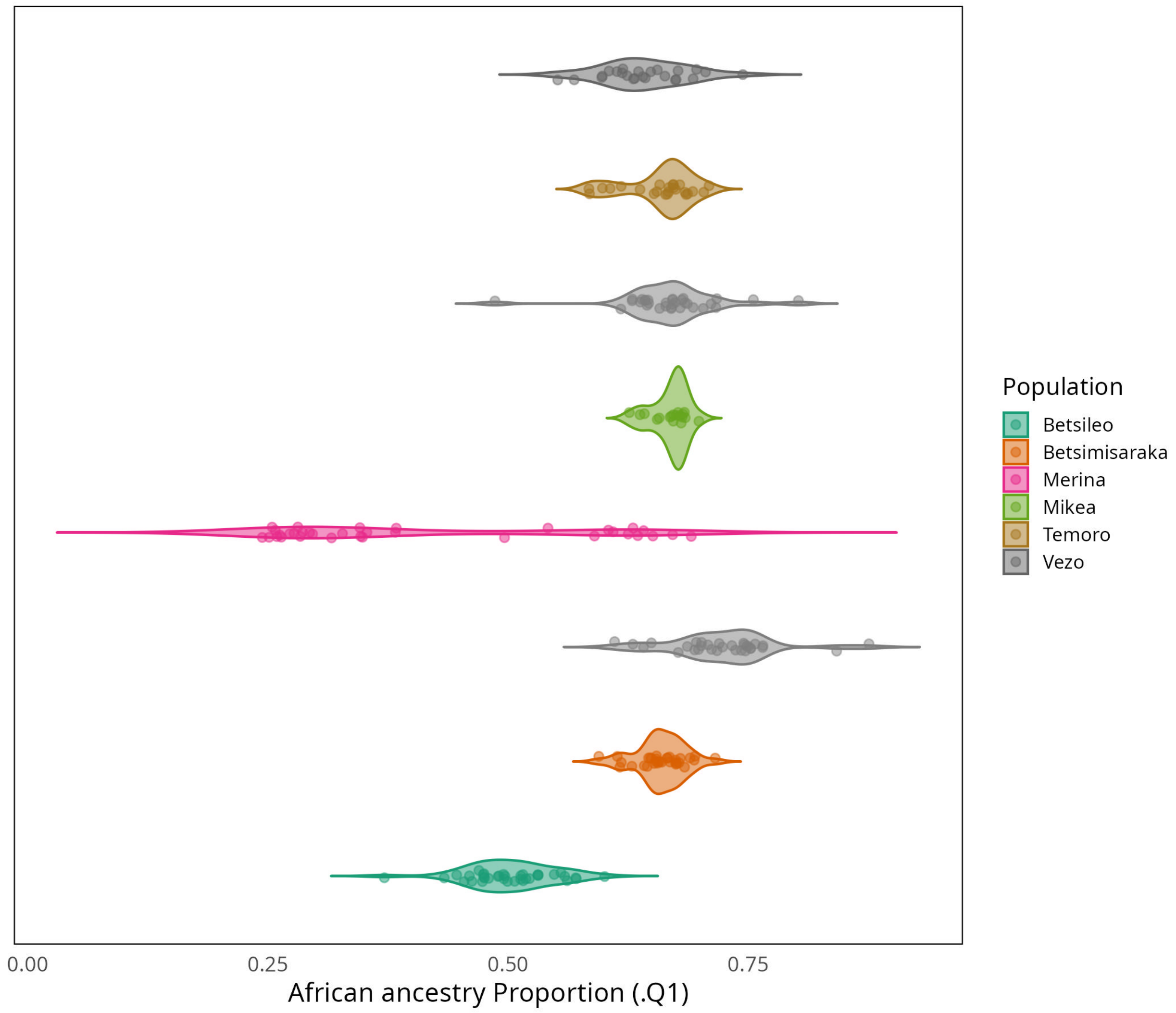


Figure 5: African ancestry proportion for each Malagasy population derived from supervised admixture at K = 2.

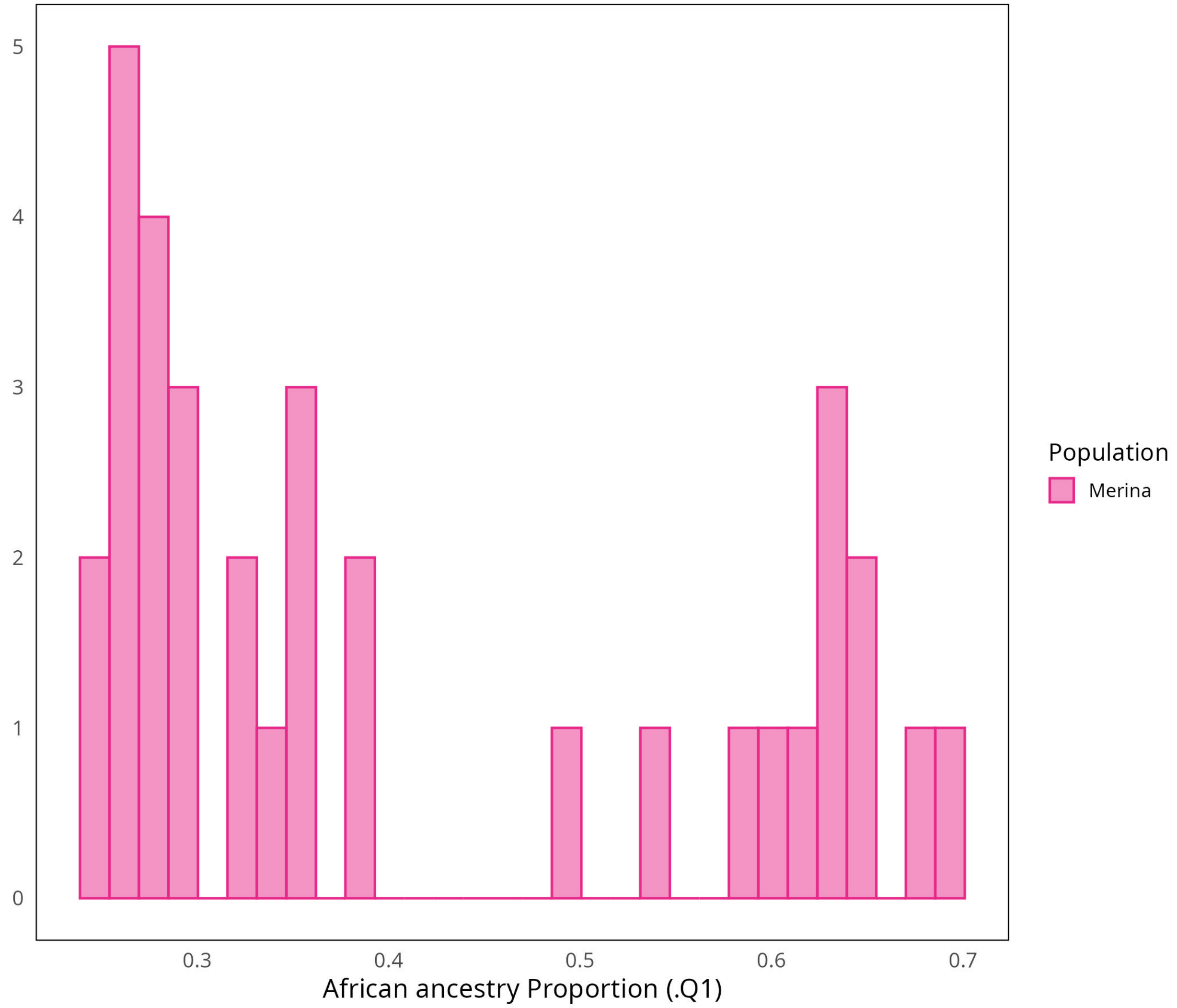


Figure 6: African ancestry proportion for Merina individuals derived from supervised admixture at K = 2, showing the bimodal distribution of ancestry among Merina individuals..

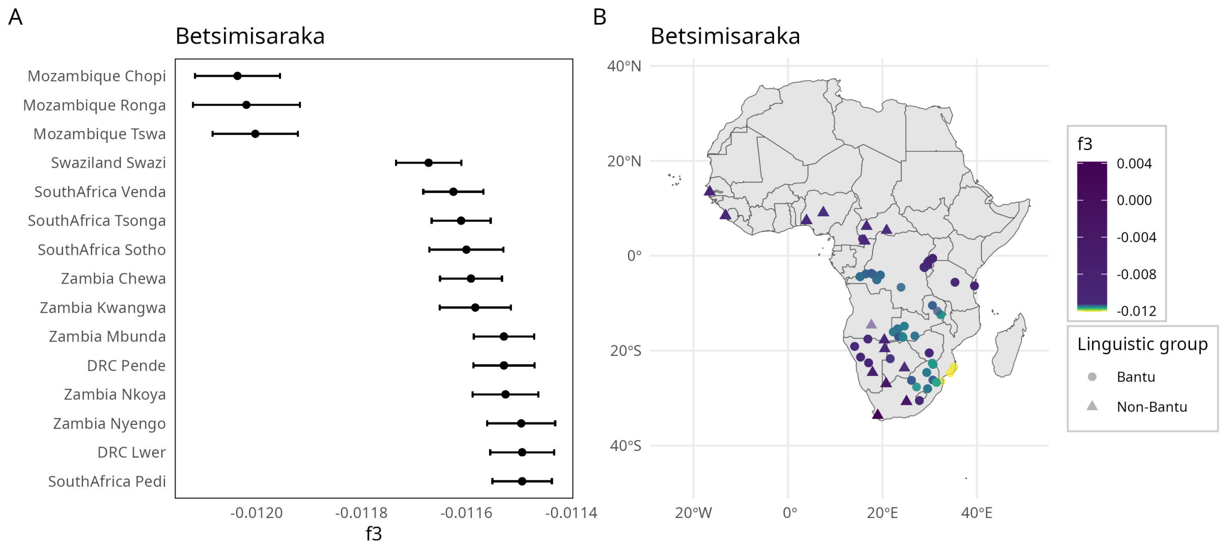


Figure 7: A) f3 statistic values across different African references for the Betsimisaraka, showing the 15 lowest f3 estimates, with Borneo as the Asian reference. B) f3 statistic values across different African references, with Borneo as the Asian reference, plotted by location for populations where coordinates were available in metadata. Colour gradient was determined by taking the standard error of the minimum f3 value and multiplying this by 10. This scale was chosen to emphasise the very best fitting f3 values, and values outside this range are coloured dark blue.

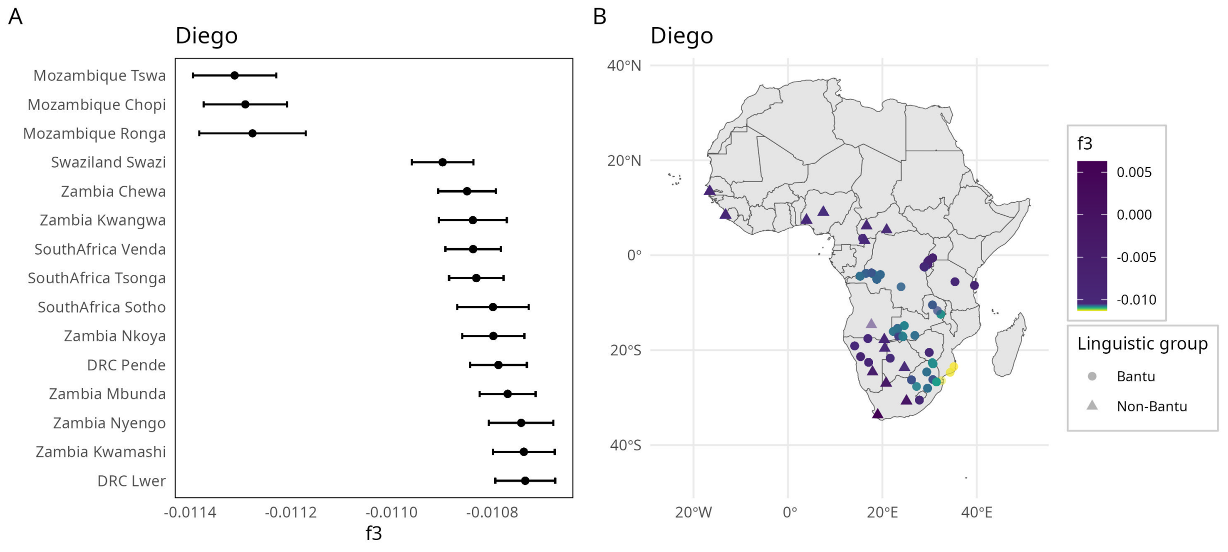


Figure 8: A) f3 statistic values across different African references for individuals from Diego, showing the 15 lowest f3 estimates, with Borneo as the Asian reference. B) f3 statistic values across different African references, with Borneo as the Asian reference, plotted by location for populations where coordinates were available in metadata. Colour gradient was determined by taking the standard error of the minimum f3 value and multiplying this by 10. This scale was chosen to emphasise the very best fitting f3 values, and values outside this range are coloured dark blue.

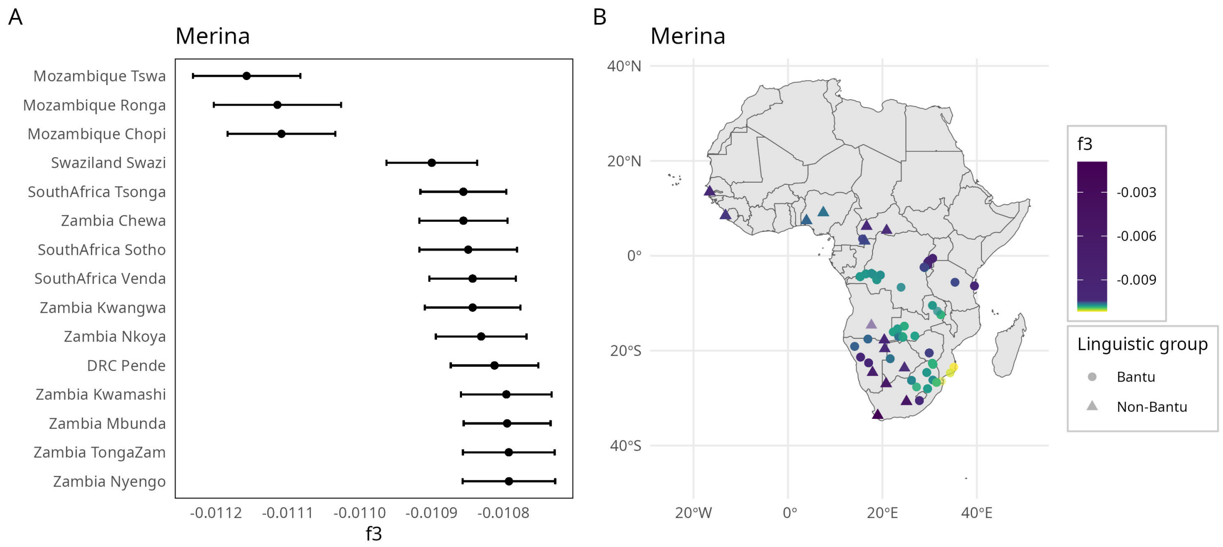


Figure 9: A) f3 statistic values across different African references for the Merina, showing the 15 lowest f3 estimates, with Borneo as the Asian reference. B) f3 statistic values across different African references, with Borneo as the Asian reference, plotted by location for populations where coordinates were available in metadata. Colour gradient was determined by taking the standard error of the minimum f3 value and multiplying this by 10. This scale was chosen to emphasise the very best fitting f3 values, and values outside this range are coloured dark blue.

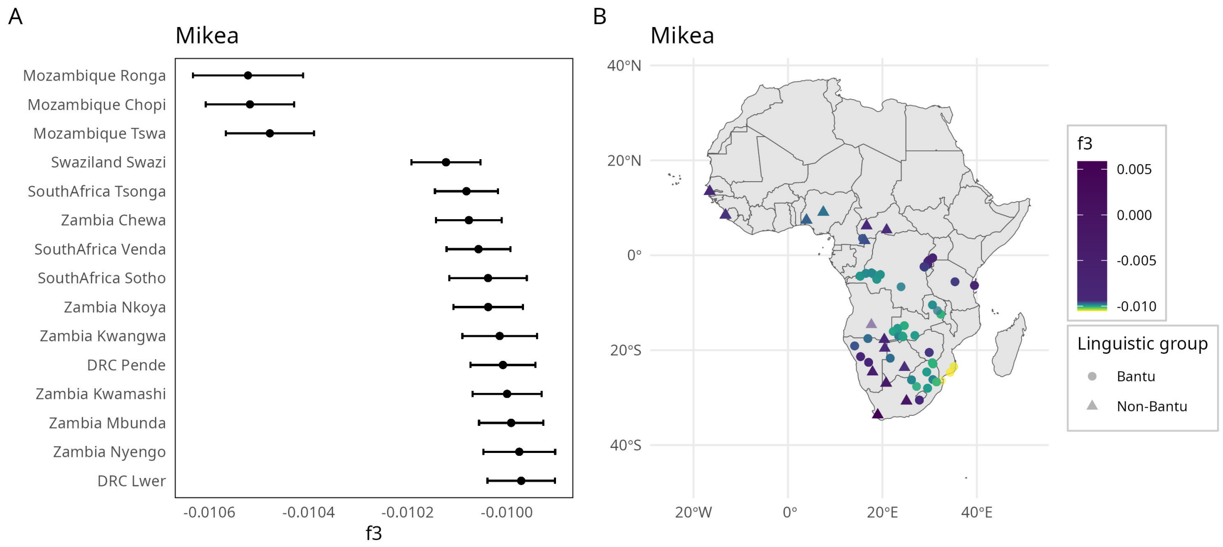


Figure 10: A) f3 statistic values across different African references for the Mikea, showing the 15 lowest f3 estimates, with Borneo as the Asian reference. B) f3 statistic values across different African references, with Borneo as the Asian reference, plotted by location for populations where coordinates were available in metadata. Colour gradient was determined by taking the standard error of the minimum f3 value and multiplying this by 10. This scale was chosen to emphasise the very best fitting f3 values, and values outside this range are coloured dark blue.

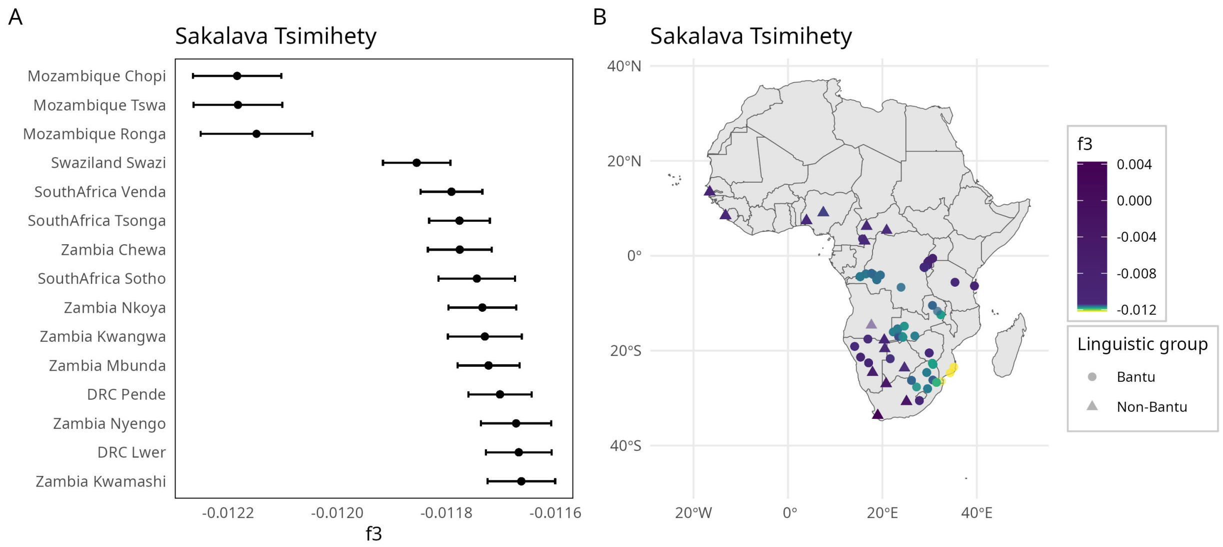


Figure 11: A) f3 statistic values across different African references for the Sakalava and Tsimihety, showing the 15 lowest f3 estimates, with Borneo as the Asian reference. B) f3 statistic values across different African references, with Borneo as the Asian reference, plotted by location for populations where coordinates were available in metadata. Colour gradient was determined by taking the standard error of the minimum f3 value and multiplying this by 10. This scale was chosen to emphasise the very best fitting f3 values, and values outside this range are coloured dark blue.

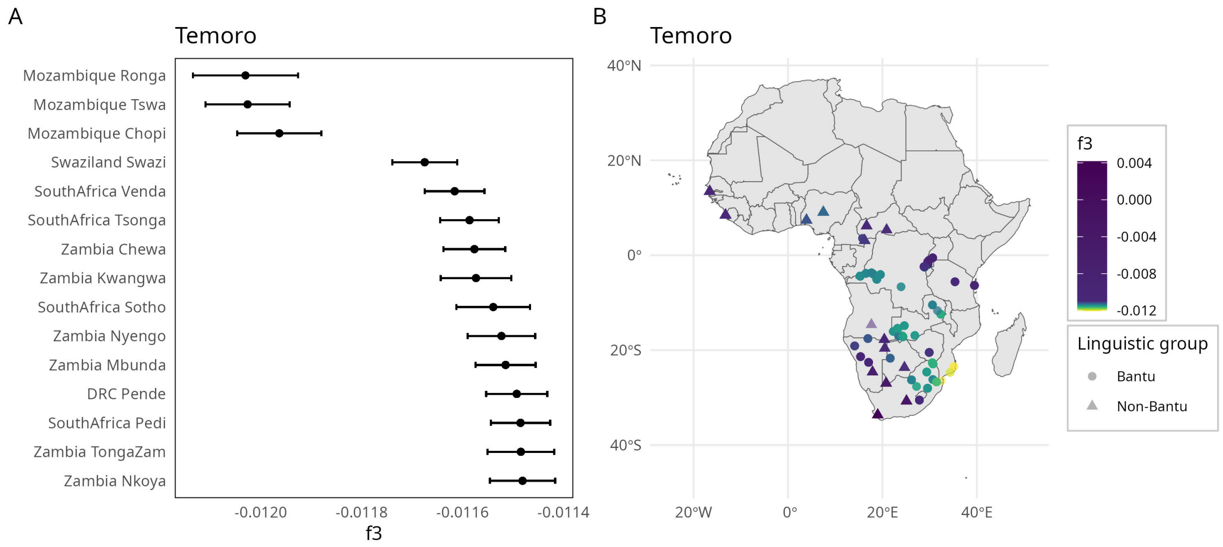


Figure 12: A) f3 statistic values across different African references for the Temoro, showing the 15 lowest f3 estimates, with Borneo as the Asian reference. B) f3 statistic values across different African references, with Borneo as the Asian reference, plotted by location for populations where coordinates were available in metadata. Colour gradient was determined by taking the standard error of the minimum f3 value and multiplying this by 10. This scale was chosen to emphasise the very best fitting f3 values, and values outside this range are coloured dark blue.

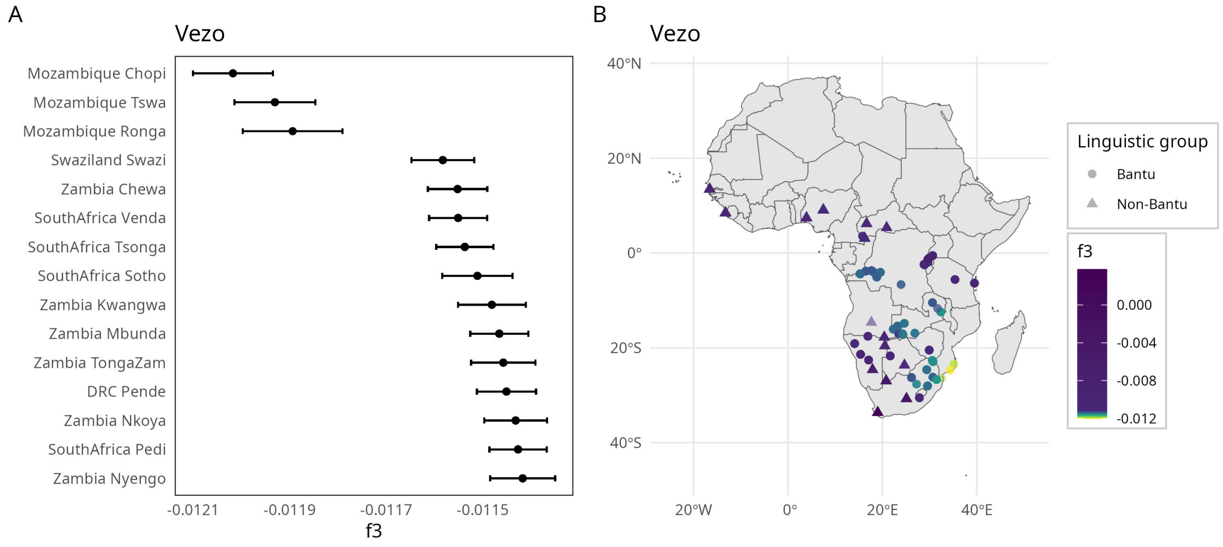


Figure 13: A) f3 statistic values across different African references for the Vezo, showing the 15 lowest f3 estimates, with Borneo as the Asian reference. B) f3 statistic values across different African references, with Borneo as the Asian reference, plotted by location for populations where coordinates were available in metadata. Colour gradient was determined by taking the standard error of the minimum f3 value and multiplying this by 10. This scale was chosen to emphasise the very best fitting f3 values, and values outside this range are coloured dark blue.

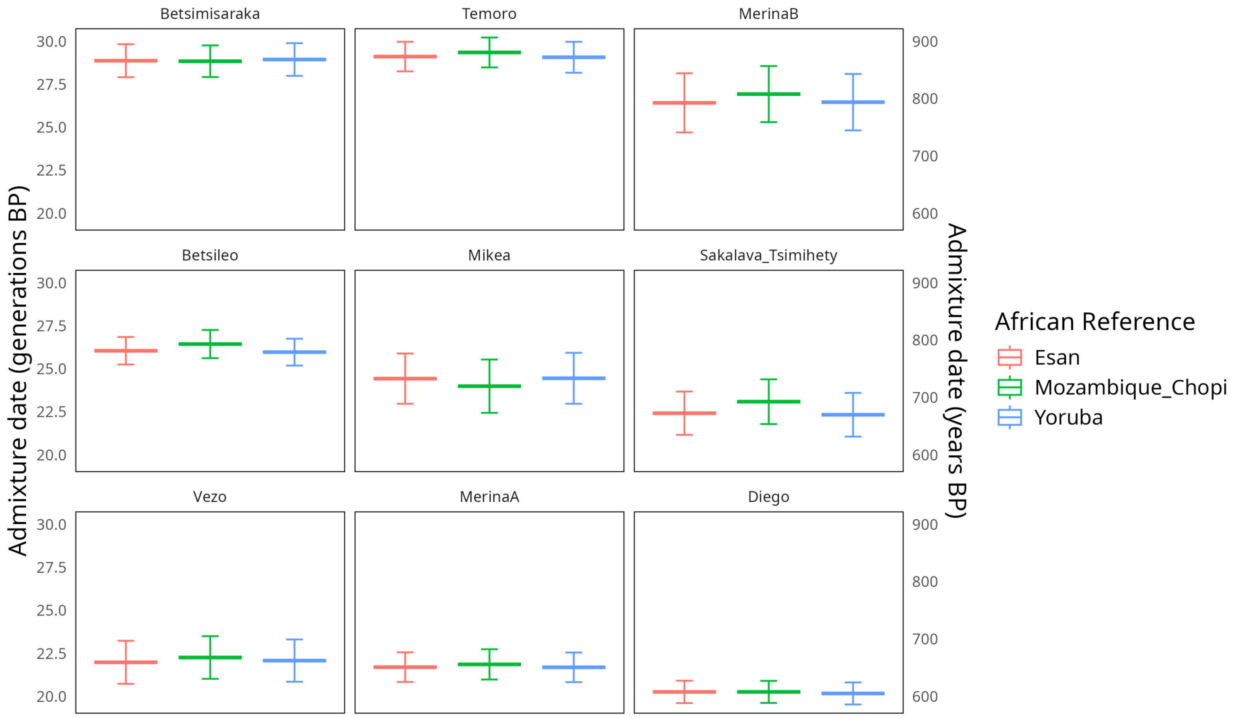


Figure 14: Estimated admixture dates for Malagasy populations using MALDER (Pickrell et al., 2014), with Borneo as an Asian source population and varied African source populations. Horizontal bar indicates estimated date of admixture, vertical bars represent standard error from leave-one-chromosome-out jackknifing.

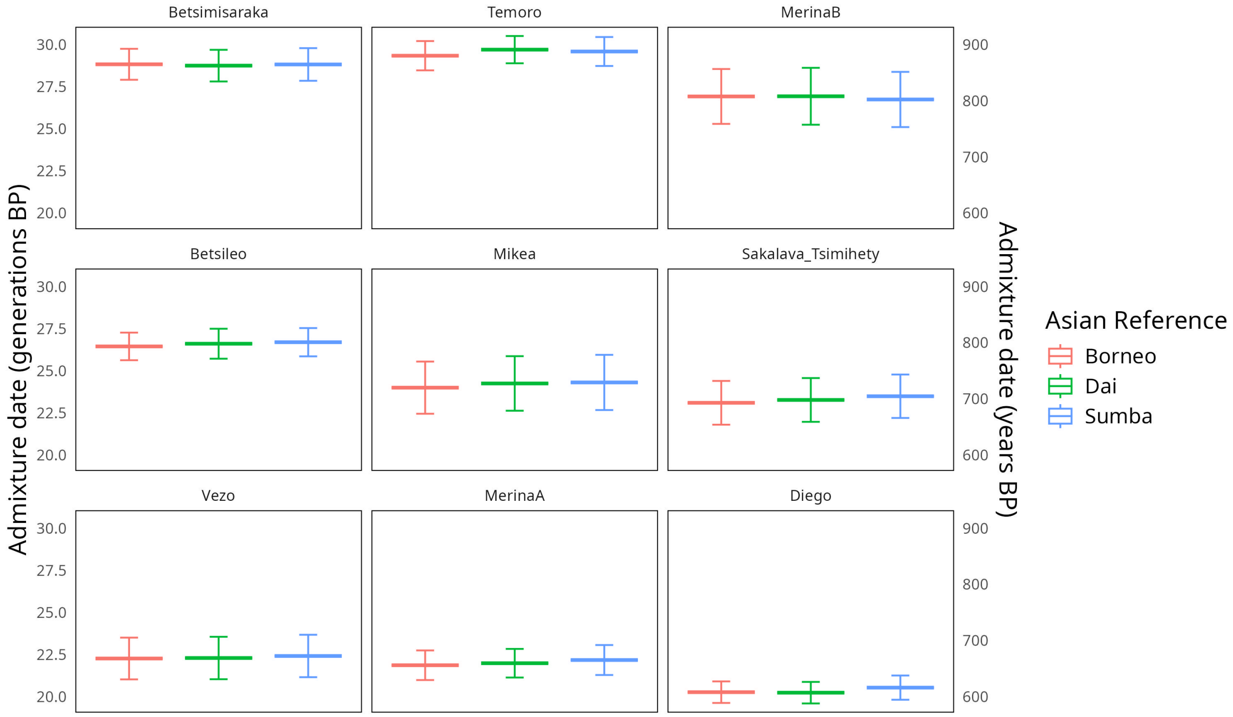


Figure 15: Estimated admixture dates for Malagasy populations using MALDER (Pickrell et al., 2014), with Mozambique as an African source population and varied Asian source populations. Horizontal bar indicates estimated date of admixture, vertical bars represent standard error from leave-one-chromosome-out jackknifing.

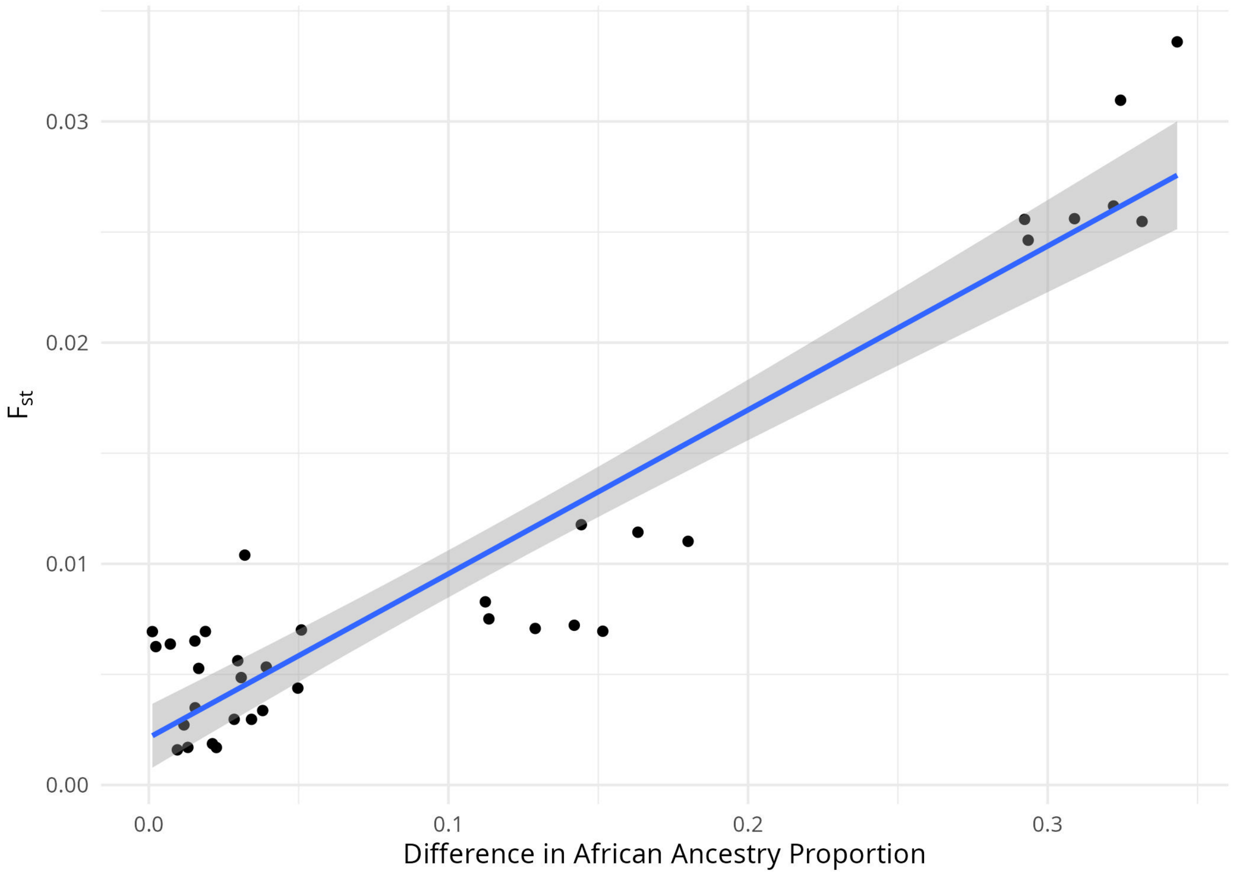


Figure 16: Difference in African ancestry proportion, as determined by f4 ratio, and Pairwise FST for pairs of Malagasy populations.

Table 1: Sample sizes for each population in the global dataset after filtering.

| Population | Sample size | source |
|---|---|---|
| Angola_Khwe | 14 | Fortes-Lima et al., 2024 |
| Angola_Sekele | 7 | Fortes-Lima et al., 2024 |
| Angola_Vasekele | 4 | Fortes-Lima et al., 2024 |
| Angola_Xun | 2 | Fortes-Lima et al., 2024 |
| Bengali, Bengal | 86 | 1000 Genomes Project Consortium, 2015 |
| Betsileo | 36 | This study |
| Betsimisaraka | 33 | This study |
| Borneo | 39 | Brucato et al., 2016 |
| Botswana_Ghanzi | 25 | Fortes-Lima et al., 2024 |
| Botswana_GuiGhanaKgal | 9 | Fortes-Lima et al., 2024 |
| Botswana_Kalanga | 3 | Fortes-Lima et al., 2024 |
| Botswana_Kgalagadi | 6 | Fortes-Lima et al., 2024 |
| Botswana_TswanaBot | 6 | Fortes-Lima et al., 2024 |
| CAR_Banda | 10 | Fortes-Lima et al., 2024 |
| CAR_DzangaShangaPeople | 16 | Fortes-Lima et al., 2024 |
| CAR_Gbaya | 21 | Fortes-Lima et al., 2024 |
| CAR_Mpiemo | 9 | Fortes-Lima et al., 2024 |
| CAR_Nzakara | 8 | Fortes-Lima et al., 2024 |
| CAR_Zande | 6 | Fortes-Lima et al., 2024 |
| Han, China | 105 | 1000 Genomes Project Consortium, 2015 |
| Diego | 31 | This study |
| DRC_Balobo | 1 | Fortes-Lima et al., 2024 |
| DRC_Bandundu | 1 | Fortes-Lima et al., 2024 |
| DRC_Bangubangu | 1 | Fortes-Lima et al., 2024 |
| DRC_BanguBangu | 2 | Fortes-Lima et al., 2024 |
| DRC_Binji | 1 | Fortes-Lima et al., 2024 |
| DRC_Binza | 1 | Fortes-Lima et al., 2024 |
| DRC_Boo | 1 | Fortes-Lima et al., 2024 |
| DRC_Ciokwe | 1 | Fortes-Lima et al., 2024 |
| DRC_Cokwe | 1 | Fortes-Lima et al., 2024 |
| DRC_Ding | 42 | Fortes-Lima et al., 2024 |
| DRC_Fang | 1 | Fortes-Lima et al., 2024 |
| DRC_Haavu | 1 | Fortes-Lima et al., 2024 |
| DRC_Hendo | 1 | Fortes-Lima et al., 2024 |
| DRC_Hungan | 3 | Fortes-Lima et al., 2024 |
| DRC_Kango | 1 | Fortes-Lima et al., 2024 |
| DRC_Kanyoka | 1 | Fortes-Lima et al., 2024 |
| DRC_Kuba | 3 | Fortes-Lima et al., 2024 |
| DRC_Kusu | 1 | Fortes-Lima et al., 2024 |
| DRC_Lele | 5 | Fortes-Lima et al., 2024 |
| DRC_Lika | 1 | Fortes-Lima et al., 2024 |
| DRC_Luba | 1 | Fortes-Lima et al., 2024 |
| DRC_LubaKasai | 2 | Fortes-Lima et al., 2024 |
| DRC_LubaKatanga | 2 | Fortes-Lima et al., 2024 |
| DRC_LubaLulua | 6 | Fortes-Lima et al., 2024 |

Table 1: Sample sizes for each population in the global dataset after filtering. *(continued)*

| Population | Sample size | source |
|---|---|---|
| DRC_Lulua | 1 | Fortes-Lima et al., 2024 |
| DRC_Lunda | 2 | Fortes-Lima et al., 2024 |
| DRC_Luozi | 2 | Fortes-Lima et al., 2024 |
| DRC_Lwer | 17 | Fortes-Lima et al., 2024 |
| DRC_Mangbetu | 1 | Fortes-Lima et al., 2024 |
| DRC_Manyanga | 35 | Fortes-Lima et al., 2024 |
| DRC_Mbaama | 1 | Fortes-Lima et al., 2024 |
| DRC_Mbala | 20 | Fortes-Lima et al., 2024 |
| DRC_Mbudza | 2 | Fortes-Lima et al., 2024 |
| DRC_Mbunda | 1 | Fortes-Lima et al., 2024 |
| DRC_Mbuun | 31 | Fortes-Lima et al., 2024 |
| DRC_MbweneNgong | 1 | Fortes-Lima et al., 2024 |
| DRC_Mongo | 1 | Fortes-Lima et al., 2024 |
| DRC_MongoNkundo | 8 | Fortes-Lima et al., 2024 |
| DRC_Mpe | 4 | Fortes-Lima et al., 2024 |
| DRC_Mpiin | 2 | Fortes-Lima et al., 2024 |
| DRC_Mpur | 1 | Fortes-Lima et al., 2024 |
| DRC_Myene | 1 | Fortes-Lima et al., 2024 |
| DRC_Nande | 1 | Fortes-Lima et al., 2024 |
| DRC_Nandi | 2 | Fortes-Lima et al., 2024 |
| DRC_Ndibu | 7 | Fortes-Lima et al., 2024 |
| DRC_Ngbaka | 1 | Fortes-Lima et al., 2024 |
| DRC_Ngbandi | 1 | Fortes-Lima et al., 2024 |
| DRC_Ngombe | 2 | Fortes-Lima et al., 2024 |
| DRC_Ngong | 4 | Fortes-Lima et al., 2024 |
| DRC_Ngwi | 67 | Fortes-Lima et al., 2024 |
| DRC_Nienge | 1 | Fortes-Lima et al., 2024 |
| DRC_Nsambaan | 1 | Fortes-Lima et al., 2024 |
| DRC_Nsong | 2 | Fortes-Lima et al., 2024 |
| DRC_Ntandu | 4 | Fortes-Lima et al., 2024 |
| DRC_NunuBobangi | 3 | Fortes-Lima et al., 2024 |
| DRC_NunuMushie | 1 | Fortes-Lima et al., 2024 |
| DRC_Nzadi | 7 | Fortes-Lima et al., 2024 |
| DRC_Nzebi | 1 | Fortes-Lima et al., 2024 |
| DRC_Pende | 21 | Fortes-Lima et al., 2024 |
| DRC_Rega | 12 | Fortes-Lima et al., 2024 |
| DRC_RundiHutu | 1 | Fortes-Lima et al., 2024 |
| DRC_Ruund | 1 | Fortes-Lima et al., 2024 |
| DRC_Sakata | 3 | Fortes-Lima et al., 2024 |
| DRC_Sengele | 1 | Fortes-Lima et al., 2024 |
| DRC_Shi | 129 | Fortes-Lima et al., 2024 |
| DRC_Songe | 3 | Fortes-Lima et al., 2024 |
| DRC_Teke | 1 | Fortes-Lima et al., 2024 |
| DRC_Tetela | 3 | Fortes-Lima et al., 2024 |
| DRC_Wonk | 4 | Fortes-Lima et al., 2024 |

Table 1: Sample sizes for each population in the global dataset after filtering. *(continued)*

| Population | Sample size | source |
|---|---|---|
| DRC_Yaka | 5 | Fortes-Lima et al., 2024 |
| DRC_Yans | 27 | Fortes-Lima et al., 2024 |
| DRC_Yombe | 9 | Fortes-Lima et al., 2024 |
| DRC_Zimba | 1 | Fortes-Lima et al., 2024 |
| Esan, Nigeria | 98 | 1000 Genomes Project Consortium, 2015 |
| Mandinka, The Gambia | 113 | 1000 Genomes Project Consortium, 2015 |
| Kinh, Vietnam | 90 | 1000 Genomes Project Consortium, 2015 |
| Lesotho_SothoSouthern | 7 | Fortes-Lima et al., 2024 |
| Luhye, Kenya | 94 | 1000 Genomes Project Consortium, 2015 |
| Merina | 34 | This study |
| Mikea | 18 | Pierron et al., 2014 |
| Mozambique_Chopi | 4 | Fortes-Lima et al., 2024 |
| Mozambique_Chuwabu | 3 | Fortes-Lima et al., 2024 |
| Mozambique_Makhuwa | 1 | Fortes-Lima et al., 2024 |
| Mozambique_Nhungwe | 3 | Fortes-Lima et al., 2024 |
| Mozambique_Ronga | 2 | Fortes-Lima et al., 2024 |
| Mozambique_Sena | 1 | Fortes-Lima et al., 2024 |
| Mozambique_TongaMoz | 3 | Fortes-Lima et al., 2024 |
| Mozambique_Tsonga | 7 | Fortes-Lima et al., 2024 |
| Mozambique_Tswa | 4 | Fortes-Lima et al., 2024 |
| Mende, Sierra Leone | 84 | 1000 Genomes Project Consortium, 2015 |
| Namibia_Damara | 21 | Fortes-Lima et al., 2024 |
| Namibia_Herero | 23 | Fortes-Lima et al., 2024 |
| Namibia_Himba | 22 | Fortes-Lima et al., 2024 |
| Namibia_Kgalagadi | 8 | Fortes-Lima et al., 2024 |
| Namibia_Nama | 15 | Fortes-Lima et al., 2024 |
| Namibia_TsumkweKung | 10 | Fortes-Lima et al., 2024 |
| Namibia_Wambo | 22 | Fortes-Lima et al., 2024 |
| Punjabi, Pakistan | 96 | 1000 Genomes Project Consortium, 2015 |
| Rwanda_Nkore | 18 | Fortes-Lima et al., 2024 |
| Sakalava_Tsimihety | 32 | This study |
| SouthAfrica_Bhaca | 12 | Fortes-Lima et al., 2024 |
| SouthAfrica_Coloured | 11 | Fortes-Lima et al., 2024 |
| SouthAfrica_ColouredColesberg | 12 | Fortes-Lima et al., 2024 |
| SouthAfrica_ColouredWellington | 19 | Fortes-Lima et al., 2024 |
| SouthAfrica_Hlubi | 2 | Fortes-Lima et al., 2024 |
| SouthAfrica_Karretjie | 9 | Fortes-Lima et al., 2024 |
| SouthAfrica_Khomani | 28 | Fortes-Lima et al., 2024 |
| SouthAfrica_Mputhi | 3 | Fortes-Lima et al., 2024 |
| SouthAfrica_Ndebele | 8 | Fortes-Lima et al., 2024 |
| SouthAfrica_Pedi | 28 | Fortes-Lima et al., 2024 |
| SouthAfrica_Shona | 1 | Fortes-Lima et al., 2024 |
| SouthAfrica_Sotho | 7 | Fortes-Lima et al., 2024 |
| SouthAfrica_SothoNorthern | 20 | Fortes-Lima et al., 2024 |
| SouthAfrica_SothoSouthern | 25 | Fortes-Lima et al., 2024 |

Table 1: Sample sizes for each population in the global dataset after filtering. *(continued)*

| Population | Sample size | source |
|---|---|---|
| SouthAfrica_Soto | 1 | Fortes-Lima et al., 2024 |
| SouthAfrica_Swazi | 5 | Fortes-Lima et al., 2024 |
| SouthAfrica_Tsonga | 26 | Fortes-Lima et al., 2024 |
| SouthAfrica_Tswana | 32 | Fortes-Lima et al., 2024 |
| SouthAfrica_Venda | 22 | Fortes-Lima et al., 2024 |
| SouthAfrica_Xhosa | 30 | Fortes-Lima et al., 2024 |
| SouthAfrica_Zulu | 52 | Fortes-Lima et al., 2024 |
| Sumba | 29 | Cox et al., 2016 |
| Swaziland_Swazi | 14 | Fortes-Lima et al., 2024 |
| Tanzania_Mixed | 1 | Fortes-Lima et al., 2024 |
| Tanzania_TanzaniaMixed | 14 | Fortes-Lima et al., 2024 |
| Temoro | 24 | Pierron et al., 2014 |
| Timor | 7 | Cox et al., 2016 |
| Uganda_Fumbira | 15 | Fortes-Lima et al., 2024 |
| Uganda_Kiga | 15 | Fortes-Lima et al., 2024 |
| Uganda_Konzo | 5 | Fortes-Lima et al., 2024 |
| Uganda_Nkore | 18 | Fortes-Lima et al., 2024 |
| Vezo | 24 | Pierron et al., 2014 |
| Yoruba, Nigeria | 104 | 1000 Genomes Project Consortium, 2015 |
| Zambia_Bemba | 11 | Fortes-Lima et al., 2024 |
| Zambia_Chewa | 17 | Fortes-Lima et al., 2024 |
| Zambia_Fwe | 14 | Fortes-Lima et al., 2024 |
| Zambia_Kalanga | 12 | Fortes-Lima et al., 2024 |
| Zambia_Kaonde | 4 | Fortes-Lima et al., 2024 |
| Zambia_Kavango | 1 | Fortes-Lima et al., 2024 |
| Zambia_Kgalagadi | 10 | Fortes-Lima et al., 2024 |
| Zambia_Kwamashi | 16 | Fortes-Lima et al., 2024 |
| Zambia_Kwangari | 1 | Fortes-Lima et al., 2024 |
| Zambia_Kwangwa | 8 | Fortes-Lima et al., 2024 |
| Zambia_Lozi | 23 | Fortes-Lima et al., 2024 |
| Zambia_Luyana | 5 | Fortes-Lima et al., 2024 |
| Zambia_Mbukushu | 8 | Fortes-Lima et al., 2024 |
| Zambia_Mbunda | 25 | Fortes-Lima et al., 2024 |
| Zambia_Ngoni | 7 | Fortes-Lima et al., 2024 |
| Zambia_Nkoya | 14 | Fortes-Lima et al., 2024 |
| Zambia_Nsenga | 4 | Fortes-Lima et al., 2024 |
| Zambia_Nyengo | 10 | Fortes-Lima et al., 2024 |
| Zambia_Shanjo | 9 | Fortes-Lima et al., 2024 |
| Zambia_Shona | 1 | Fortes-Lima et al., 2024 |
| Zambia_ShonaZam | 4 | Fortes-Lima et al., 2024 |
| Zambia_TongaZam | 10 | Fortes-Lima et al., 2024 |
| Zambia_Tswana | 2 | Fortes-Lima et al., 2024 |
| Zambia_Tumbuka | 4 | Fortes-Lima et al., 2024 |
| Zanzibar_Swahili | 22 | Fortes-Lima et al., 2024 |
| Zimbabwe_Remba | 32 | Fortes-Lima et al., 2024 |

Table 1: Sample sizes for each population in the global dataset after filtering. *(continued)*

| Population | Sample size | source |
|---|---|---|
| Total | 2744 | Total |

Table 2: Summary of analyses and software used in this study.

| Analysis | Package/function | Dataset | Individuals Removed | n indivs | Loci filtering | n SNPs |
|---|---|---|---|---|---|---|
| Malagasy PCA | tidypopgen: gt__pca__randomSVD | Malagasy dataset | Missingness > 0.1; KING kinship > 0.0884 | Non-autosomal; Missingness < 0.05; HWE p > 0.01; MAF > 0.025; LD clumping | 233 | 227410 |
| Global PCA | tidypopgen: gt__pca__randomSVD | Global dataset | Missingness > 0.1; KING kinship > 0.0884 | Non-autosomal; Missingness < 0.05; HWE p > 0.01; MAF > 0.025; LD clumping | 2744 | 198196 |
| Unsupervised clustering | tidypopgen: gt__admixture | Global dataset | Missingness > 0.1; KING kinship > 0.0884 | Non-autosomal; Missingness < 0.05; HWE p > 0.01; MAF > 0.025; LD clumping | 2744 | 198196 |
| Supervised clustering | ADMIXTURE: ADMIXTURE | Subset of Global dataset | Missingness > 0.1; KING kinship > 0.0884 | Non-autosomal; Missingness < 0.05; HWE p > 0.01; MAF > 0.025; LD clumping | 299 | 198196 |
| F3 statistics | tidypopgen: gt__extract__f2; admixtools2 (version 2.0.8) (Maier and Patterson, 2024): qp3pop | Global dataset | Missingness > 0.1; KING kinship > 0.0884 | Non-autosomal; Missingness < 0.05; HWE p > 0.01; MAF > 0.025; LD clumping | 2744 | 198196 |
| F4 ratio statistics | tidypopgen: gt__extract__f2; admixtools2 (version 2.0.8) (Maier and Patterson, 2024): qpf4ratio | Global dataset + Pan and European outgroups | Missingness > 0.1; KING kinship > 0.0884 | Non-autosomal; Missingness < 0.05; HWE p > 0.01; MAF > 0.025; LD clumping | 2792 | 180431 |
| MALDER | MALDER (version 1.0) (Pickrell et al., 2014): MALDER | Global dataset | Missingness > 0.1; KING kinship > 0.0884 | Non-autosomal; Missingness < 0.05; HWE p > 0.01; MAF > 0.025 | 2744 | 425904 |
| Pairwise FST | tidypopgen: pairwise__pop__fst - method = "WC84" (Weir and Cockerham, 1984) | Global dataset | Missingness > 0.1; KING kinship > 0.0884 | Non-autosomal; Missingness < 0.05; HWE p > 0.01; MAF > 0.025; LD clumping | 2744 | 198196 |

Table 3: Minimum f3 statistic values across all combinations of references for each Malagasy population.

| Malagasy | Asian | African | est | se | z | p |
|---|---|---|---|---|---|---|
| Betsileo | Borneo | Mozambique_Ronga | -0.0123638 | 0.0000925 | -133.67820 | 0 |
| Betsimisaraka | Borneo | Mozambique_Chopi | -0.0120364 | 0.0000810 | -148.58346 | 0 |
| Diego | Borneo | Mozambique_Tswa | -0.0113118 | 0.0000828 | -136.57844 | 0 |
| Merina | Borneo | Mozambique_Tswa | -0.0111576 | 0.0000747 | -149.26685 | 0 |
| Mikea | Borneo | Mozambique_Ronga | -0.0105223 | 0.0001111 | -94.68672 | 0 |
| Sakalava Tsimihety | Borneo | Mozambique_Chopi | -0.0121836 | 0.0000811 | -150.27787 | 0 |
| Temoro | Borneo | Mozambique_Ronga | -0.0120305 | 0.0001042 | -115.42556 | 0 |
| Vezo | Borneo | Mozambique_Chopi | -0.0120179 | 0.0000829 | -144.89537 | 0 |

Table 4: Minimum z-scores for f3 statistics across all combinations for each Malagasy population.

| Malagasy | Asian | African | est | se | z | p |
|---|---|---|---|---|---|---|
| Betsileo | Borneo | DRC_Shi | -0.0115276 | 5.51e-05 | -209.2158 | 0 |
| Betsimisaraka | Borneo | DRC_Shi | -0.0110031 | 5.16e-05 | -213.0884 | 0 |
| Diego | Borneo | DRC_Shi | -0.0102260 | 4.85e-05 | -210.7807 | 0 |
| Merina | Borneo | DRC_Shi | -0.0104804 | 5.67e-05 | -184.9265 | 0 |
| Mikea | Borneo | DRC_Ngwi | -0.0099074 | 6.14e-05 | -161.3010 | 0 |
| Sakalava Tsimihety | Borneo | DRC_Shi | -0.0111734 | 5.08e-05 | -220.1194 | 0 |
| Temoro | Borneo | DRC_Shi | -0.0110101 | 5.37e-05 | -205.2034 | 0 |
| Vezo | Borneo | DRC_Shi | -0.0109646 | 5.46e-05 | -200.6620 | 0 |

Table 5: Table 5 is provided in an accompanying Excel file.

| Detailed Results (See accompanying Excel file) |
|---|

Table 6: f4 ratios calculated with admixtools. f4 ratios were calculated in the configurations f4(GBR,Chimpanzee;Malagasy,Borneo) / f4(GBR,Chimpanzee;Esan,Borneo) and f4(GBR,Chimpanzee;Malagasy,Esan) / f4(GBR,Chimpanzee;Borneo,Esan)

| pop1 | pop2 | pop3 | pop4 | pop5 | alpha | se | z |
|---|---|---|---|---|---|---|---|
| GBR | chimp_ref | Betsimisaraka | Borneo | ESN | 0.6087948 | 0.0056688 | 107.39458 |
| GBR | chimp_ref | MerinaA | Borneo | ESN | 0.2773051 | 0.0074150 | 37.39799 |
| GBR | chimp_ref | MerinaB | Borneo | ESN | 0.5695955 | 0.0082825 | 68.77100 |
| GBR | chimp_ref | Diego | Borneo | ESN | 0.6205166 | 0.0056107 | 110.59508 |
| GBR | chimp_ref | Sakalava_Tsimihety | Borneo | ESN | 0.5992745 | 0.0054699 | 109.55800 |
| GBR | chimp_ref | Betsileo | Borneo | ESN | 0.4572821 | 0.0059435 | 76.93771 |
| GBR | chimp_ref | Temoro | Borneo | ESN | 0.5862443 | 0.0060979 | 96.13881 |
| GBR | chimp_ref | Vezo | Borneo | ESN | 0.5707852 | 0.0062701 | 91.03236 |
| GBR | chimp_ref | Mikea | Borneo | ESN | 0.6016209 | 0.0071687 | 83.92276 |

Table 7: MALDER results for all Malagasy populations, with varied African and Asian source populations. p value: significance of model tested. test pop: Malagasy target population. ref A: Asian population. ref B: African population. 2 ref z score: z score. 2 ref decay: Date, in generations, since admixture began with standard error. 2 ref amp exp: Amplitude of the LD decay curve. amp aff: Affine term estimating population substructure.

| p_value | test_pop | ref A | ref B | 2-ref z-score | 2-ref decay | 2-ref amp_exp | amp_aff |
|---|---|---|---|---|---|---|---|
| 4.6e-222 | Betsileo | Borneo | Esan | 31.81 | 26.05 +/- 0.80 | 0.00216033 +/- 0.00006791 | 0.0000299 |
| 3.6e-234 | Betsileo | Borneo | Yoruba | 32.67 | 25.97 +/- 0.78 | 0.00214609 +/- 0.00006568 | 0.0000296 |
| 5.7e-230 | Betsileo | Borneo | Mozambique_Chopi | 32.38 | 26.44 +/- 0.82 | 0.00227784 +/- 0.00006829 | 0.0000310 |
| 3.5e-188 | Betsileo | Sumba | Esan | 29.26 | 26.26 +/- 0.82 | 0.00191612 +/- 0.00006549 | 0.0000261 |
| 1.7e-196 | Betsileo | Sumba | Yoruba | 29.91 | 26.18 +/- 0.80 | 0.00190193 +/- 0.00006360 | 0.0000258 |
| 1.5e-200 | Betsileo | Sumba | Mozambique_Chopi | 30.22 | 26.69 +/- 0.84 | 0.00202856 +/- 0.00006714 | 0.0000271 |
| 8.6e-200 | Betsileo | Dai | Esan | 30.16 | 26.21 +/- 0.87 | 0.00216652 +/- 0.00006781 | 0.0000304 |
| 1.5e-205 | Betsileo | Dai | Yoruba | 30.59 | 26.14 +/- 0.85 | 0.00215206 +/- 0.00006551 | 0.0000300 |
| 1.3e-197 | Betsileo | Dai | Mozambique_Chopi | 29.99 | 26.60 +/- 0.89 | 0.00228532 +/- 0.00006899 | 0.0000315 |
| 1.1e-196 | Betsimisaraka | Borneo | Esan | 29.92 | 28.87 +/- 0.96 | 0.00187297 +/- 0.00004607 | 0.0000083 |
| 7.2e-204 | Betsimisaraka | Borneo | Yoruba | 30.47 | 28.94 +/- 0.95 | 0.00186972 +/- 0.00004541 | 0.0000081 |
| 6.9e-217 | Betsimisaraka | Borneo | Mozambique_Chopi | 31.43 | 28.84 +/- 0.92 | 0.00197274 +/- 0.00004997 | 0.0000086 |
| 4.5e-185 | Betsimisaraka | Sumba | Esan | 29.01 | 28.90 +/- 1.00 | 0.00165692 +/- 0.00005122 | 0.0000072 |
| 2.2e-189 | Betsimisaraka | Sumba | Yoruba | 29.35 | 28.96 +/- 0.99 | 0.00165296 +/- 0.00005075 | 0.0000070 |
| 1.6e-193 | Betsimisaraka | Sumba | Mozambique_Chopi | 29.68 | 28.83 +/- 0.97 | 0.00175133 +/- 0.00005520 | 0.0000074 |
| 1.3e-199 | Betsimisaraka | Dai | Esan | 30.14 | 28.81 +/- 0.96 | 0.00186998 +/- 0.00004787 | 0.0000080 |
| 6.6e-205 | Betsimisaraka | Dai | Yoruba | 30.54 | 28.88 +/- 0.95 | 0.00186641 +/- 0.00004697 | 0.0000078 |
| 3.4e-204 | Betsimisaraka | Dai | Mozambique_Chopi | 30.49 | 28.76 +/- 0.94 | 0.00196995 +/- 0.00005151 | 0.0000083 |
| 1.5e-144 | Diego | Borneo | Esan | 25.60 | 20.26 +/- 0.65 | 0.00146813 +/- 0.00005735 | 0.0000493 |
| 4.1e-148 | Diego | Borneo | Yoruba | 25.92 | 20.17 +/- 0.64 | 0.00145881 +/- 0.00005628 | 0.0000494 |
| 4.4e-159 | Diego | Borneo | Mozambique_Chopi | 26.87 | 20.26 +/- 0.64 | 0.00153534 +/- 0.00005713 | 0.0000514 |
| 2.2e-116 | Diego | Sumba | Esan | 22.93 | 20.57 +/- 0.73 | 0.00131657 +/- 0.00005741 | 0.0000432 |
| 9.8e-120 | Diego | Sumba | Yoruba | 23.27 | 20.46 +/- 0.71 | 0.00130741 +/- 0.00005619 | 0.0000433 |
| 6.7e-124 | Diego | Sumba | Mozambique_Chopi | 23.67 | 20.53 +/- 0.72 | 0.00137919 +/- 0.00005826 | 0.0000452 |
| 4.7e-130 | Diego | Dai | Esan | 24.26 | 20.24 +/- 0.66 | 0.00146933 +/- 0.00006055 | 0.0000496 |
| 1.1e-133 | Diego | Dai | Yoruba | 24.60 | 20.14 +/- 0.64 | 0.00145999 +/- 0.00005934 | 0.0000497 |
| 2.3e-138 | Diego | Dai | Mozambique_Chopi | 25.04 | 20.23 +/- 0.64 | 0.00153750 +/- 0.00006140 | 0.0000517 |
| 1.2e-228 | Merina | Borneo | Esan | 32.28 | 22.65 +/- 0.70 | 0.00176093 +/- 0.00005003 | 0.0004215 |
| 5.5e-236 | Merina | Borneo | Yoruba | 32.80 | 22.66 +/- 0.69 | 0.00175499 +/- 0.00004855 | 0.0004206 |
| 1.5e-226 | Merina | Borneo | Mozambique_Chopi | 32.13 | 23.18 +/- 0.72 | 0.00186187 +/- 0.00004419 | 0.0004401 |
| 1.1e-227 | Merina | Sumba | Esan | 32.21 | 22.68 +/- 0.65 | 0.00154560 +/- 0.00004798 | 0.0003678 |
| 2.2e-238 | Merina | Sumba | Yoruba | 32.97 | 22.68 +/- 0.64 | 0.00153947 +/- 0.00004669 | 0.0003670 |
| 1.6e-250 | Merina | Sumba | Mozambique_Chopi | 33.81 | 23.32 +/- 0.69 | 0.00164251 +/- 0.00004452 | 0.0003852 |
| 3.8e-233 | Merina | Dai | Esan | 32.60 | 22.74 +/- 0.70 | 0.00176475 +/- 0.00005140 | 0.0004208 |

Table 7: MALDER results for all Malagasy populations, with varied African and Asian source populations. p value: significance of model tested. test pop: Malagasy target population. ref A: Asian population. ref B: African population. 2 ref z score: z score. 2 ref decay: Date, in generations, since admixture began with standard error. 2 ref amp exp: Amplitude of the LD decay curve. amp aff: Affine term estimating population substructure. *(continued)*

| p_value | test_pop | ref A | ref B | 2-ref z-score | 2-ref decay | 2-ref amp_exp | amp_aff |
|---|---|---|---|---|---|---|---|
| 3.9e-244 | Merina | Dai | Yoruba | 33.37 | 22.75 +/- 0.68 | 0.00175932 +/- 0.00004984 | 0.0004199 |
| 1.8e-241 | Merina | Dai | Mozambique_Chopi | 33.19 | 23.29 +/- 0.70 | 0.00186752 +/- 0.00004541 | 0.0004395 |
| 5.5e-141 | MerinaA | Borneo | Esan | 25.28 | 21.70 +/- 0.86 | 0.00172501 +/- 0.00006674 | 0.0000510 |
| 2.5e-139 | MerinaA | Borneo | Yoruba | 25.13 | 21.69 +/- 0.86 | 0.00172051 +/- 0.00006620 | 0.0000510 |
| 1.8e-135 | MerinaA | Borneo | Mozambique_Chopi | 24.77 | 21.86 +/- 0.88 | 0.00180736 +/- 0.00006513 | 0.0000546 |
| 1.8e-128 | MerinaA | Sumba | Esan | 24.11 | 21.96 +/- 0.86 | 0.00151669 +/- 0.00006290 | 0.0000432 |
| 4.2e-129 | MerinaA | Sumba | Yoruba | 24.17 | 21.94 +/- 0.86 | 0.00151185 +/- 0.00006254 | 0.0000433 |
| 8.1e-137 | MerinaA | Sumba | Mozambique_Chopi | 24.90 | 22.17 +/- 0.89 | 0.00159416 +/- 0.00006327 | 0.0000466 |
| 1.4e-149 | MerinaA | Dai | Esan | 26.05 | 21.80 +/- 0.84 | 0.00173212 +/- 0.00006550 | 0.0000519 |
| 1.3e-149 | MerinaA | Dai | Yoruba | 26.05 | 21.80 +/- 0.84 | 0.00172795 +/- 0.00006479 | 0.0000519 |
| 1.2e-146 | MerinaA | Dai | Mozambique_Chopi | 25.79 | 21.98 +/- 0.85 | 0.00181659 +/- 0.00006355 | 0.0000556 |
| 2.3e-53 | MerinaB | Borneo | Esan | 15.38 | 26.42 +/- 1.72 | 0.00182359 +/- 0.00006434 | 0.0000235 |
| 1.2e-58 | MerinaB | Borneo | Yoruba | 16.15 | 26.46 +/- 1.64 | 0.00181557 +/- 0.00006173 | 0.0000232 |
| 4.2e-61 | MerinaB | Borneo | Mozambique_Chopi | 16.49 | 26.93 +/- 1.63 | 0.00193439 +/- 0.00005846 | 0.0000234 |
| 5.1e-52 | MerinaB | Sumba | Esan | 15.18 | 26.11 +/- 1.72 | 0.00159222 +/- 0.00006515 | 0.0000201 |
| 8.6e-57 | MerinaB | Sumba | Yoruba | 15.88 | 26.13 +/- 1.65 | 0.00158453 +/- 0.00006284 | 0.0000198 |
| 8.8e-60 | MerinaB | Sumba | Mozambique_Chopi | 16.31 | 26.75 +/- 1.64 | 0.00170122 +/- 0.00005956 | 0.0000200 |
| 1.9e-49 | MerinaB | Dai | Esan | 14.78 | 26.44 +/- 1.79 | 0.00181330 +/- 0.00006328 | 0.0000236 |
| 4.9e-54 | MerinaB | Dai | Yoruba | 15.48 | 26.48 +/- 1.71 | 0.00180611 +/- 0.00006077 | 0.0000233 |
| 3.8e-57 | MerinaB | Dai | Mozambique_Chopi | 15.93 | 26.94 +/- 1.69 | 0.00192369 +/- 0.00005553 | 0.0000236 |
| 6.2e-63 | Mikea | Borneo | Esan | 16.74 | 24.43 +/- 1.46 | 0.00179737 +/- 0.00007812 | 0.0000009 |
| 4.2e-61 | Mikea | Borneo | Yoruba | 16.49 | 24.45 +/- 1.48 | 0.00179523 +/- 0.00007745 | 0.0000008 |
| 8.6e-54 | Mikea | Borneo | Mozambique_Chopi | 15.44 | 23.99 +/- 1.55 | 0.00184531 +/- 0.00007972 | 0.0000014 |
| 6.7e-59 | Mikea | Sumba | Esan | 16.18 | 24.75 +/- 1.53 | 0.00158879 +/- 0.00007863 | 0.0000010 |
| 1.3e-56 | Mikea | Sumba | Yoruba | 15.86 | 24.77 +/- 1.56 | 0.00158630 +/- 0.00007822 | 0.0000009 |
| 6.3e-50 | Mikea | Sumba | Mozambique_Chopi | 14.86 | 24.30 +/- 1.64 | 0.00163606 +/- 0.00008013 | 0.0000015 |
| 2.2e-58 | Mikea | Dai | Esan | 16.11 | 24.68 +/- 1.53 | 0.00181233 +/- 0.00008078 | 0.0000010 |
| 1.1e-56 | Mikea | Dai | Yoruba | 15.87 | 24.70 +/- 1.56 | 0.00181012 +/- 0.00008012 | 0.0000009 |
| 2.4e-50 | Mikea | Dai | Mozambique_Chopi | 14.92 | 24.24 +/- 1.62 | 0.00186109 +/- 0.00008254 | 0.0000015 |
| 1.1e-70 | Sakalava_Tsimihety | Borneo | Esan | 17.77 | 22.42 +/- 1.26 | 0.00177727 +/- 0.00007455 | 0.0000396 |
| 4.3e-69 | Sakalava_Tsimihety | Borneo | Yoruba | 17.57 | 22.33 +/- 1.27 | 0.00176631 +/- 0.00007391 | 0.0000395 |
| 3.3e-70 | Sakalava_Tsimihety | Borneo | Mozambique_Chopi | 17.71 | 23.09 +/- 1.30 | 0.00188537 +/- 0.00007484 | 0.0000422 |
| 1.4e-73 | Sakalava_Tsimihety | Sumba | Esan | 18.14 | 22.70 +/- 1.25 | 0.00157199 +/- 0.00007657 | 0.0000344 |
| 3.5e-71 | Sakalava_Tsimihety | Sumba | Yoruba | 17.84 | 22.60 +/- 1.27 | 0.00156087 +/- 0.00007598 | 0.0000343 |

Table 7: MALDER results for all Malagasy populations, with varied African and Asian source populations. p value: significance of model tested. test pop: Malagasy target population. ref A: Asian population. ref B: African population. 2 ref z score: z score. 2 ref decay: Date, in generations, since admixture began with standard error. 2 ref amp exp: Amplitude of the LD decay curve. amp aff: Affine term estimating population substructure. *(continued)*

| p_value | test_pop | ref A | ref B | 2-ref z-score | 2-ref decay | 2-ref amp_exp | amp_aff |
|---|---|---|---|---|---|---|---|
| 1.2e-73 | Sakalava_Tsimihety | Sumba | Mozambique_Chopi | 18.15 | 23.48 +/- 1.29 | 0.00167853 +/- 0.00007755 | 0.0000367 |
| 1.4e-72 | Sakalava_Tsimihety | Dai | Esan | 18.02 | 22.60 +/- 1.25 | 0.00177662 +/- 0.00007776 | 0.0000398 |
| 3.9e-71 | Sakalava_Tsimihety | Dai | Yoruba | 17.83 | 22.51 +/- 1.26 | 0.00176524 +/- 0.00007692 | 0.0000396 |
| 5.3e-72 | Sakalava_Tsimihety | Dai | Mozambique_Chopi | 17.94 | 23.26 +/- 1.30 | 0.00188434 +/- 0.00007879 | 0.0000423 |
| 9.6e-133 | Temoro | Borneo | Esan | 24.52 | 29.11 +/- 0.86 | 0.00194337 +/- 0.00007926 | 0.0000182 |
| 1e-128 | Temoro | Borneo | Yoruba | 24.14 | 29.07 +/- 0.90 | 0.00193347 +/- 0.00008011 | 0.0000178 |
| 7.4e-147 | Temoro | Borneo | Mozambique_Chopi | 25.81 | 29.35 +/- 0.87 | 0.00205756 +/- 0.00007973 | 0.0000186 |
| 5.3e-104 | Temoro | Sumba | Esan | 21.66 | 29.34 +/- 0.87 | 0.00172607 +/- 0.00007970 | 0.0000165 |
| 2.7e-101 | Temoro | Sumba | Yoruba | 21.37 | 29.28 +/- 0.90 | 0.00171615 +/- 0.00008032 | 0.0000162 |
| 7.5e-112 | Temoro | Sumba | Mozambique_Chopi | 22.47 | 29.60 +/- 0.86 | 0.00183343 +/- 0.00008158 | 0.0000169 |
| 5.4e-130 | Temoro | Dai | Esan | 24.26 | 29.48 +/- 0.82 | 0.00195981 +/- 0.00008079 | 0.0000181 |
| 6.1e-127 | Temoro | Dai | Yoruba | 23.97 | 29.42 +/- 0.85 | 0.00194883 +/- 0.00008131 | 0.0000178 |
| 2.8e-142 | Temoro | Dai | Mozambique_Chopi | 25.40 | 29.71 +/- 0.81 | 0.00207391 +/- 0.00008166 | 0.0000185 |
| 1.9e-69 | Vezo | Borneo | Esan | 17.62 | 21.98 +/- 1.25 | 0.00184944 +/- 0.00006413 | 0.0000285 |
| 5.5e-72 | Vezo | Borneo | Yoruba | 17.94 | 22.08 +/- 1.23 | 0.00184557 +/- 0.00006245 | 0.0000284 |
| 1.2e-72 | Vezo | Borneo | Mozambique_Chopi | 18.03 | 22.26 +/- 1.24 | 0.00194431 +/- 0.00005711 | 0.0000301 |
| 2.4e-67 | Vezo | Sumba | Esan | 17.34 | 22.16 +/- 1.28 | 0.00163475 +/- 0.00005995 | 0.0000246 |
| 1.6e-70 | Vezo | Sumba | Yoruba | 17.76 | 22.26 +/- 1.25 | 0.00163029 +/- 0.00005825 | 0.0000245 |
| 1.3e-70 | Vezo | Sumba | Mozambique_Chopi | 17.76 | 22.41 +/- 1.26 | 0.00172420 +/- 0.00005415 | 0.0000261 |
| 4.2e-66 | Vezo | Dai | Esan | 17.17 | 22.04 +/- 1.28 | 0.00185290 +/- 0.00006451 | 0.0000280 |
| 1.1e-68 | Vezo | Dai | Yoruba | 17.51 | 22.13 +/- 1.26 | 0.00184873 +/- 0.00006279 | 0.0000278 |
| 3.5e-70 | Vezo | Dai | Mozambique_Chopi | 17.71 | 22.29 +/- 1.26 | 0.00194813 +/- 0.00005747 | 0.0000296 |

Table 8: Pairwise FST values between all Malagasy populations.

| | Betsileo | Betsimisaraka | Diego.Mixed | Merina.A | Merina.B | Mikea | Sakalava.Tsimihety | Temoro | Vezo |
|---|---|---|---|---|---|---|---|---|---|
| Betsileo | NA | 0.0069522 | 0.0114288 | 0.0110161 | 0.0082805 | 0.0117686 | 0.0072215 | 0.0070768 | 0.0075091 |
| Betsimisaraka | 0.0069522 | NA | 0.0027162 | 0.0254780 | 0.0053308 | 0.0063718 | 0.0015886 | 0.0016943 | 0.0033650 |
| Diego Mixed | 0.0114288 | 0.0027162 | NA | 0.0335991 | 0.0070089 | 0.0069401 | 0.0018644 | 0.0029681 | 0.0043794 |
| Merina A | 0.0110161 | 0.0254780 | 0.0335991 | NA | 0.0255723 | 0.0309598 | 0.0261779 | 0.0256050 | 0.0246308 |
| Merina B | 0.0082805 | 0.0053308 | 0.0070089 | 0.0255723 | NA | 0.0103931 | 0.0056181 | 0.0052702 | 0.0069354 |
| Mikea | 0.0117686 | 0.0063718 | 0.0069401 | 0.0309598 | 0.0103931 | NA | 0.0062564 | 0.0065102 | 0.0048565 |
| Sakalava/Tsimihety | 0.0072215 | 0.0015886 | 0.0018644 | 0.0261779 | 0.0056181 | 0.0062564 | NA | 0.0016993 | 0.0029689 |
| Temoro | 0.0070768 | 0.0016943 | 0.0029681 | 0.0256050 | 0.0052702 | 0.0065102 | 0.0016993 | NA | 0.0034909 |
| Vezo | 0.0075091 | 0.0033650 | 0.0043794 | 0.0246308 | 0.0069354 | 0.0048565 | 0.0029689 | 0.0034909 | NA |

Kent, W. James, Charles W. Sugnet, Terrence S. Furey, Krishna M. Roskin, Tom H. Pringle, Alan M. Zahler, and and David Haussler. 2002. “The Human Genome Browser at UCSC.” *Genome Research* 12 (6): 996–1006. https://doi.org/10.1101/gr.229102.

Manichaikul, Ani, Josyf C. Mychaleckyj, Stephen S. Rich, Kathy Daly, Michèle Sale, and Wei-Min Chen. 2010. “Robust Relationship Inference in Genome-Wide Association Studies.” *Bioinformatics* 26 (22): 2867–73. https://doi.org/10.1093/bioinformatics/btq559.

Privé, Florian, Hugues Aschard, Andrey Ziyatdinov, and Michael G B Blum. 2018. “Efficient Analysis of Large-Scale Genome-Wide Data with Two R Packages: Bigstatsr and Bigsnpr.” *Bioinformatics* 34 (16): 2781–87. https://doi.org/10.1093/bioinformatics/bty185.
